# Hippocampal representations of distance, space, and direction and their plasticity predict navigational performance

**DOI:** 10.1101/2020.12.30.424858

**Authors:** Jason J Moore, Jesse D Cushman, Lavanya Acharya, Mayank R Mehta

## Abstract

The hippocampus is implicated in episodic memory and allocentric spatial navigation. However, spatial selectivity is insufficient to navigate; one needs information about the distance and direction to the reward on a specific journey. The nature of these representations, whether they are expressed in an episodic-like sequence, and their relationship with navigational performance are unknown. We recorded single units from dorsal CA1 of the hippocampus while rats navigated to an unmarked reward zone defined solely by distal visual cues, similar to the classic water maze. The allocentric spatial selectivity was substantially weaker than in typical real world tasks, despite excellent navigational performance. Instead, the majority of cells encoded path distance from the start of trials. Cells also encoded the rat’s allocentric position and head angle. Often the same cells multiplexed and encoded path distance, head direction and allocentric position in a sequence, thus encoding a journey-specific episode. The strength of neural activity and tuning strongly correlated with performance, with a temporal relationship indicating neural responses influencing behavior and vice versa. Consistent with computational models of associative Hebbian learning, neural responses showed increasing clustering and became better predictors of behaviorally relevant variables, with neurometric curves exceeding and converging to psychometric curves. These findings demonstrate that hippocampal neurons multiplex and exhibit highly plastic, task- and experience-dependent tuning to path-centric and allocentric variables to form an episode, which could mediate navigation.

## Introduction: Place cells, navigation and synaptic plasticity

The rodent hippocampus is implicated in spatial navigation using distal visual landmarks, as measured by the classic Morris water maze navigation task^1–3^. This task is thought to be mediated by cognitive mapping^4,5^, or path integration^6,7^, built from the allocentric selectivity of place cells^8^. While indirect evidence for this relationship exists^9–11^, a precise link is lacking between computational models of NMDAR-dependent synaptic plasticity^12,13^, emergent place field plasticity^13–18^, and navigational performance^1–3^. Additionally, extensive studies in primates and humans show that the hippocampus is involved in episodic memory^19–21^. Episodic-like responses have also been reported in the rodent hippocampus^22–27^. But the link between simultaneous representations of episodic and allocentric spatial signals with spatial navigation has remained unclear in the rodent hippocampus, particularly the episodic ordering of different spatial signals.

Further, these different forms of learning are hypothesized to be mediated by Hebbian plasticity, which has two components: associative learning^28,29^ and temporally-asymmetric learning in the form of spike-timing dependent plasticity (STDP)^30,31^. Computational models of learning via associative plasticity predict clustering of neural responses^32^, and navigational learning via STDP predict that place cells should increase firing rates and show anticipatory shift^12–15^. This has been observed on narrow, one dimensional paths^13–18^ but not in two dimensions, and the link between these models and navigation has remained unclear.

## Performance in virtual navigation task

Hence, we trained rats to execute a virtual navigation task similar to the Morris water maze^33^ and measured hippocampal neural responses and their experience-dependent plasticity. The use of VR maintains a visual cue-based navigational demand while entirely removing nonspecific cues, matching one of the fundamental features of cognitive mapping and the Morris water maze while removing human intervention. Combined with the appetitive nature of the task, which is similar to most place cell recording tasks, this allows the rat to run many trials in a single session, providing an ideal approach to testing computational models of navigational learning, and removing stress experienced in the water maze that impairs synaptic plasticity^34^. The trial based structure of the task allows us to explore the neural encoding of sequences of behaviorally relevant events and measures, such as initiation and direction of movement, distance travelled and the expected position of the reward.

As previously demonstrated^33^, rats readily learned to perform the virtual navigation task using only distal visual cues. In addition to latency to reward, here we measured performance as the number of rewards the rats obtained per meter (RPM) of distance traveled, similar to the path-length measure commonly used in the water maze task^1^. Paths were relatively direct in most trials, but different from each other from the four start positions to the hidden goal location (Fig. 1a), suggesting the use of a navigational strategy rather than a stereotyped motor response. To further ensure the use of a navigational strategy, experimental conditions such as the number of start positions, reward zone size, and cues on the walls were changed every 2-4 days. These changes defined “session blocks” in which to track behavioral improvement under identical conditions (Fig. 1b). Across sessions within session blocks, well-trained rats continued to improve on subsequent days (Fig. 1b), further supporting the use of a navigational strategy that is refined under changing experimental conditions. Furthermore, paths originating from different start positions were distinct from each other, even in the more difficult task conditions with eight start positions rather than four (Extended Data Fig. 1). Other behavioral measures such as time spent in the target quadrant, time spent near the reward zone, and running speed near the reward zone also demonstrated efficient navigational behavior (Extended Data Fig. 2). These measures were not significantly different in 8-start compared to 4-start tasks (Extended Data Fig. 2).

**Fig. 1.**
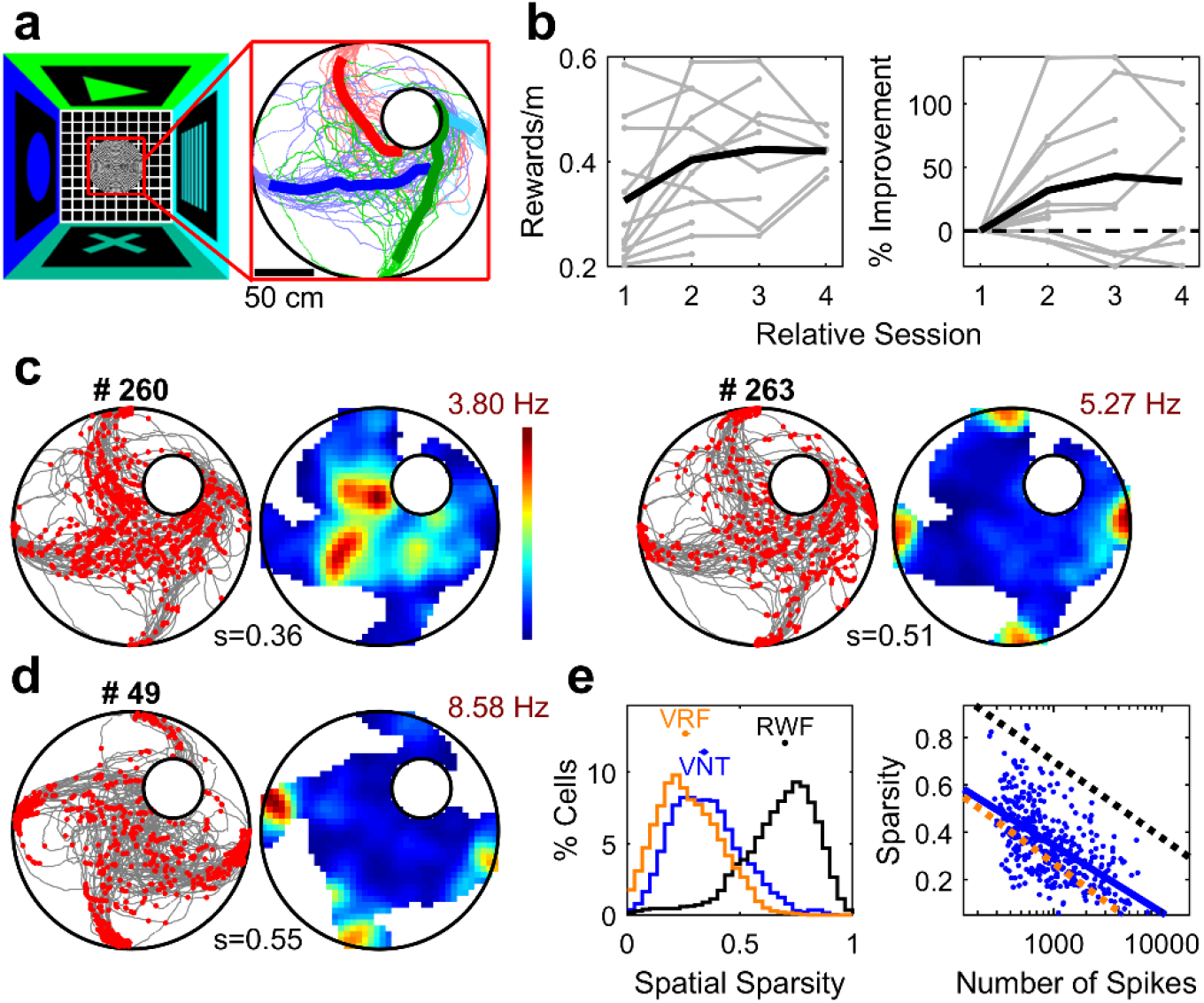
Good performance but impaired spatial selectivity in a virtual navigation task. **a,** Left, overhead schematic of virtual environment, with a 200 cm-diameter table centered in a 400×400 cm room with distinct visual cues on each wall. Right, paths from individual trials (thin lines) and mean paths from each start position (thick lines), color-coded by start position. The hidden reward zone is indicated by the white dot in the north-east quadrant. **b,** Performance, measured by rewards/meter, consistently improved across subsequent sessions in different session blocks (p=0.02, Wilcoxon sign-rank test on difference between % Improvement across consecutive days without a gap, n=27 differences). Thin gray lines indicate individual session blocks, with the thick black line indicating the mean (n=12 session blocks). **c,** Example spike plots and spatial rate maps for two individual units from the session in **a.** The left cell exhibits low spatial selectivity, while the right cell has higher spatial sparsity with fields near the start position of each trial. Gray lines indicate paths, and red dots indicate spikes. Spatial sparsity (*s*) is indicated below the rate maps. Firing rates range from 0 Hz to the maximum value indicated above the rate maps. **d,** Same as **c** but for a unit from a different behavioral session. **e,** Left, the spatial sparsity in the virtual navigation task (VNT, blue) (0.34, [0.32, 0.36], n=384 units), was slightly but significantly greater (p=1.5×10^-12^, Wilcoxon rank-sum test) than that in a 2-dimensional random foraging task in the same virtual reality system (VRF, orange) (0.26, [0.24, 0.28], n=421 units), and significantly less (p=1.2×10^-106^, Wilcoxon rank-sum test) than that in a real world random foraging task (RWF, black) (0.7, [0.69, 0.71], n=626 units). Right, the group differences in spatial sparsity were still significant after controlling for the inverse relationship between the number of spikes and sparsity (VNT vs VRF: p=0.39, two-way ANOVA, see Methods; VNT vs RWF: p=4.7×10^-6^, two-way ANOVA, see Methods).

## Very little allocentric selectivity

To investigate the neural mechanisms of this navigational task, we measured the activities of 384 putative CA1 pyramidal neurons from 4 rats in 34 sessions using tetrodes (see Methods). We first quantified the degree of allocentric spatial selectivity during this task. In contrast to typical random foraging tasks in a two-dimensional real world environment, CA1 neurons showed relatively little allocentric spatial selectivity in virtual navigation (Fig. 1c-e, Extended Data Fig. 3), similar to that found in a random foraging task in the same VR system^25^. Spatial sparsity was not significantly different between 4-start and 8-start versions of the task (Extended Data Fig. 3b-d), and these data were pooled together for all analyses unless noted.

How can a rat execute a navigation task with such low allocentric spatial selectivity? We hypothesize that hippocampal neurons could contain information about distance traveled and the direction of the reward^11,26,35,36^, which could be sufficient for navigation. However, all these variables are collinear, complicating estimates of their independent contributions to neural activity. Hence, we further developed the generalized linear model (GLM) technique^37^ to include these parameters as covariates. With this more sensitive analysis, few cells (~1/3) were significantly modulated by allocentric space (Fig. 2a, d, Extended Data Fig. 4), and these were significantly less stable than spatial fields in a RW foraging task^37^ (Extended Data Fig. 5). This instability was not due to parameter overfitting, as GLM-derived maps were fitted using 5-fold cross validation and were more stable than expected by chance, particularly for well-tuned cells (Extended Data Fig. 5). Again, there was no difference in spatial sparsity between 4-start and 8-start versions of the task (Extended Data Fig. 6). Interestingly, the distribution of peaks of these allocentric fields was neither uniformly distributed across the maze nor clustered near the start position^38^ but they were significantly clustered near the reward zone (Fig. 2e), where the occupancy was the highest (Extended Data Fig. 6). This is consistent with theories of learning based on associative synaptic plasticity^28,29,32^.

**Fig. 2.**
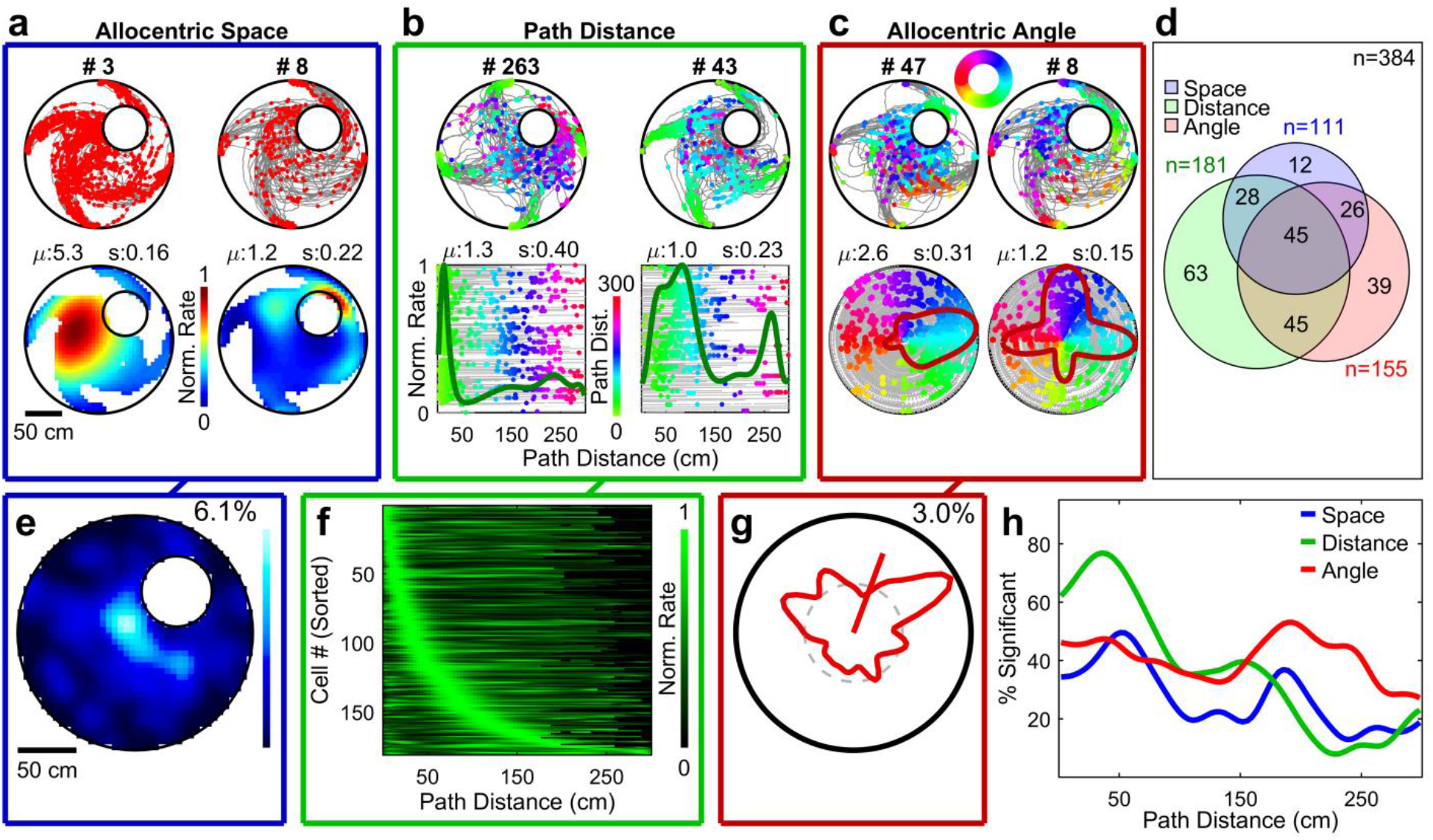
Allocentric, path-centric, and angular tuning. **a,** Spike plots (top) and GLM-derived spatial rate maps (bottom) for two separate neurons with statistically significant spatial sparsity. Color conventions are as in Fig. 1. **b,** Spike plots (top) and GLM-derived distance rate maps (green traces, bottom) for two separate neurons with statistically significant distance sparsity. Spikes in the top and bottom plots are color coded according to path distance. Elapsed time (gray) increases along the y-axis in the bottom plots. **c,** Spike plots (top) and GLM-derived angular rate maps (red traces, bottom) for two separate neurons with statistically significant angle sparsity. Spikes are color coded according to angle. In the bottom plots, session time (gray) begins at the origin and increases radially outward. **d,** 29, [24, 34]% of units were tuned for allocentric space (S). 47, [42, 52]% of units were tuned for path distance (D). 40, [35, 45]% of units were tuned for angle (A). SD (19, [15, 23]%), SA (18, [15, 23]%), DA (23, [19, 28]%), and SDA (12, [9, 15]%) represent the percentages of cells tuned for a combination of parameters. n=384 units for all percentages in **d**. **e,** Distribution of the maxima of spatial rate maps for cells significantly tuned for space (n=111). The density ranges from 0 to 6.1%. **f,** Distance rate maps for all cells significantly tuned for path distance (n=181), sorted by location of peak rate (median of 32, [24, 39] cm). **g,** Distribution of the maxima of angular rate maps for cells significantly tuned for angle (n=155). The line indicates the mean vector, at 70, [68, 72]°, and the maximum density is 3.0%. **h,** The percentage of neurons tuned for space, distance, and angle fluctuated as a function of path distance (see Methods). Spatial tuning peaked near 50 and 200 cm. Distance tuning peaked near 40 cm, and angle tuning peaked near 200 cm.

## Most neurons encode distance and angle

In contrast, nearly half of neurons were significantly modulated by path distance, defined as the distance traveled from the start of a trial, regardless of the start position (Fig. 2b, d, Extended Data Fig. 7, 8). This was true in both the 4- and 8-start tasks (Extended Data Fig. 8). These distance receptive fields were qualitatively similar to place fields in one dimensional tracks in real and virtual worlds^23^. Path distance field centers spanned approximately 200 cm but were clustered towards short distances, with more than half the cells centered before 32 cm (Fig. 2b, f). This clustering is different than many other studies that found either a more homogeneous distribution of place field centers in RW or VR linear tracks^23,38^ or a linear treadmill task^24^, or clustering near goal locations in a linear treadmill task^39^, annular water maze task^40^, or bats flying towards a visible goal^26^. This clustering mirrors the behavioral oversampling of early distances (Extended Data Fig. 8), consistent with theories of associative learning. Path distance fields often exhibited a multi-peaked structure (Extended Data Fig. 9), and were more unstable than typical place fields in the real world, but more stable than allocentric spatial maps in our experiments (Extended Data Fig. 5). This overrepresentation was not because all trials contained short distances, as distance fields were still aggregated when computed only for long trials (Extended Data Fig. 10). We found similar selectivity when trial progression was measured as time elapsed rather than distance traveled, with a small but significant preference for distance, rather than time (Extended Data Fig. 11)^41,42^. Conversely, very few cells showed clear tuning to the distance from the goal, as compared to distance from the start (Extended Data Fig. 12). Distance tuning could not be explained by motor signals alone, e.g. the act of turning at the trial beginning (Extended Data Fig. 13).

Similar to two-dimensional random foraging tasks in both RW and VR^37^, but to a greater degree, approximately 40% of CA1 neurons were significantly modulated by head angle with respect to the distal visual cues (Fig. 2c, d, g, Extended Data Fig. 14). Many neurons had multi-peaked angular tuning curves, similar to the angular maps in random foraging tasks in RW and VR (Extended Data Fig. 15)^37^. The distribution of the peak angles spanned all directions (Fig. 2c, g, Extended Data Fig. 14). However, the distribution was not homogeneous and a majority of the preferred angles clustered towards the northeast direction which, for most locations on the maze, is towards the hidden reward zone^26^. In line with the other variables, angular field density clustering matched the behavioral distribution (Extended Data Fig. 14). Stability of the angular tuning curves was slightly less than that of the distance fields (Extended Data Fig. 5), but similar to that during random foraging in VR^37^.

To determine if neurons with different codes were distinct, we quantified the overlap between the populations of cells significantly modulated by space, distance, and angle (Fig. 2d, Extended Data Fig. 16). The proportion of cells that were modulated by two or more variables was approximately equal to that expected by chance, indicating that individual CA1 neurons can multiplex and simultaneously encode head directional, allocentric, and path-centric information^43^, rather than being segregated populations^22,39,44^.

## Neural selectivity reflects behavioral performance

Are these different types of information equally available at all times, or are these three codes organized in a need-dependent way as a function the progression of a trial, i.e. an episode? We quantified the percentage of cells significantly tuned for space, distance, or angle as a function of the distance at which the cell’s firing rate was maximal (Fig. 2h, Extended Data Fig. 17). Path distance tuning was highest at short distances (<50 cm), consistent with the aggregation of distance field centers near short distances. Allocentric space tuning showed two peaks, near 50 and 200 cm, congruent with the distribution of path lengths from nearby (50cm) and distant (200cm) start locations. Angle selectivity was relatively high throughout, with a peak at 200 cm, indicating that angle tuning may be most relevant when the rat is near the reward. The relative ordering of distance, space, and then angle tuning is not expected by chance (Extended Data Fig. 17c). This indicates that each journey is encoded as an episode, which is a mixture of the three codes, each becoming more prominent when most needed.

Though all rats in this study were well trained, the performance was quite variable in each rat on a day-to-day basis (Extended Data Fig. 2d). We leveraged this to investigate the relationship between behavior and neural tuning (Fig. 3, Extended Data Fig. 18). The mean firing rate of cells in a session was significantly positively correlated with performance (Fig. 3c, Extended Data Fig. 18), as measured by rewards/meter, trial latency, trial distance, or path correlation (see Methods). This was not simply an artifact of increased running speed, as running speed was not correlated with performance (Extended Data Fig. 19). Additionally, performance was positively correlated with the percentage of cells in a given session that were significantly modulated by head angle, allocentric space, or path distance (Fig. 3c, Extended Data Fig. 18).

**Fig. 3.**
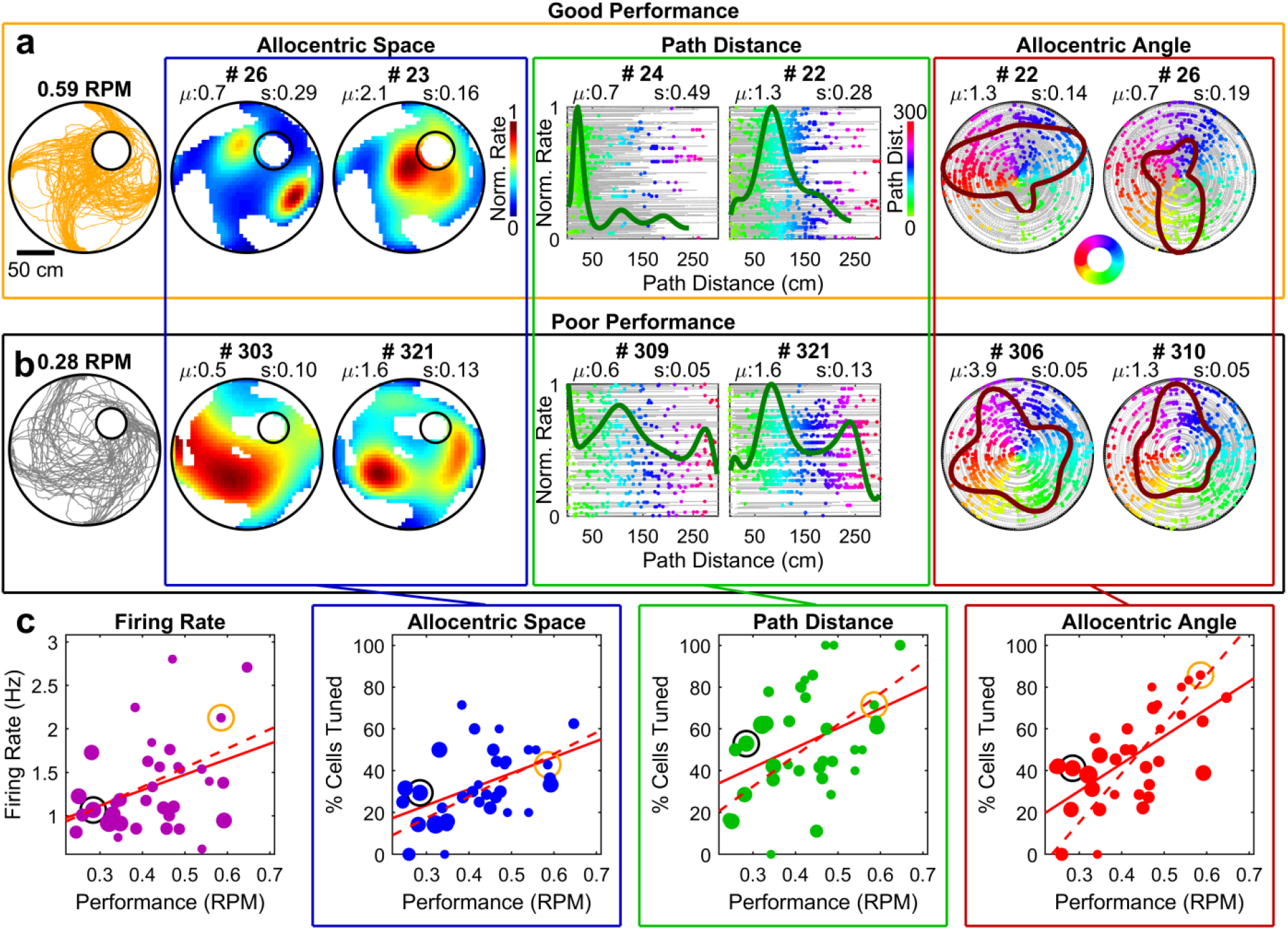
Tuning is correlated with behavior. **a,** Sample behavioral session with good behavior (RPM: Rewards per meter), with two examples each of rate maps for allocentric space, path distance, and angle, all with relatively high degrees of tuning. **b,** Sample behavioral session with poor behavior, with two examples each of rate maps for allocentric space, path distance, and angle, all with relatively low degrees of tuning. **c,** Performance was significantly positively correlated with mean firing rate (left). Each point represents a single session, with the size proportional to the number of units recorded in that session (minimum 4). The solid line is the unweighted best linear fit (R=0.36, p=0.04, two-sided *t* test for unweighted best fit, n=34 sessions here and throughout **c)** and the dashed line is the best linear fit weighted by the number of cells (R=0.33, p=0.04, see Methods). Performance was also positively correlated with the percentage of cells tuned for allocentric space (middle left, unweighted: R=0.48, p=4.4×10^-3^; weighted: R=0.48, p=5×10^-4^), the percentage of cells tuned for path distance (middle right, unweighted: R=0.40, p=0.02; weighted: R=0.41, p=5.4×10^-3^), as well as the percentage of cells tuned for angle (right, unweighted: R=0.63, p=5.8×10^-5^; weighted: R=0.56, p<1×10^-4^). The orange and black circles represent the example sessions in **a** and **b**, respectively.

## Correlated experiential plasticity of neurometric and psychometric curves within a session

There was a significant improvement in performance within behavioral sessions each day, as the trial latency and rewards per meter both improved by up to 50% as a function of trial number (Fig. 4a, Extended Data Fig. 20a), without an accompanying increase in speed (Extended Data Fig. 20b). Computational theories of navigational learning via the temporally asymmetric component of STDP or Hebbian plasticity ^12–15,30,31^ and experiments on linear tracks predict that navigational learning could be mediated by increased firing rates and predictive, or backward, shift of neural responses^13–18^. Consistently, we observed that only approximately 45% of cells were active in the initial trials of a session, and this proportion increased with trial number to 55% by trial 50, comparable to that seen in the real world ^18^ (Extended Data Fig. 20c). The mean firing rate of cells that were active also increased with trial number, from 1.5 Hz to 2.1 Hz (Fig. 4b), comparable to that seen on linear tracks in the real world ^13–18^.

**Fig. 4.**
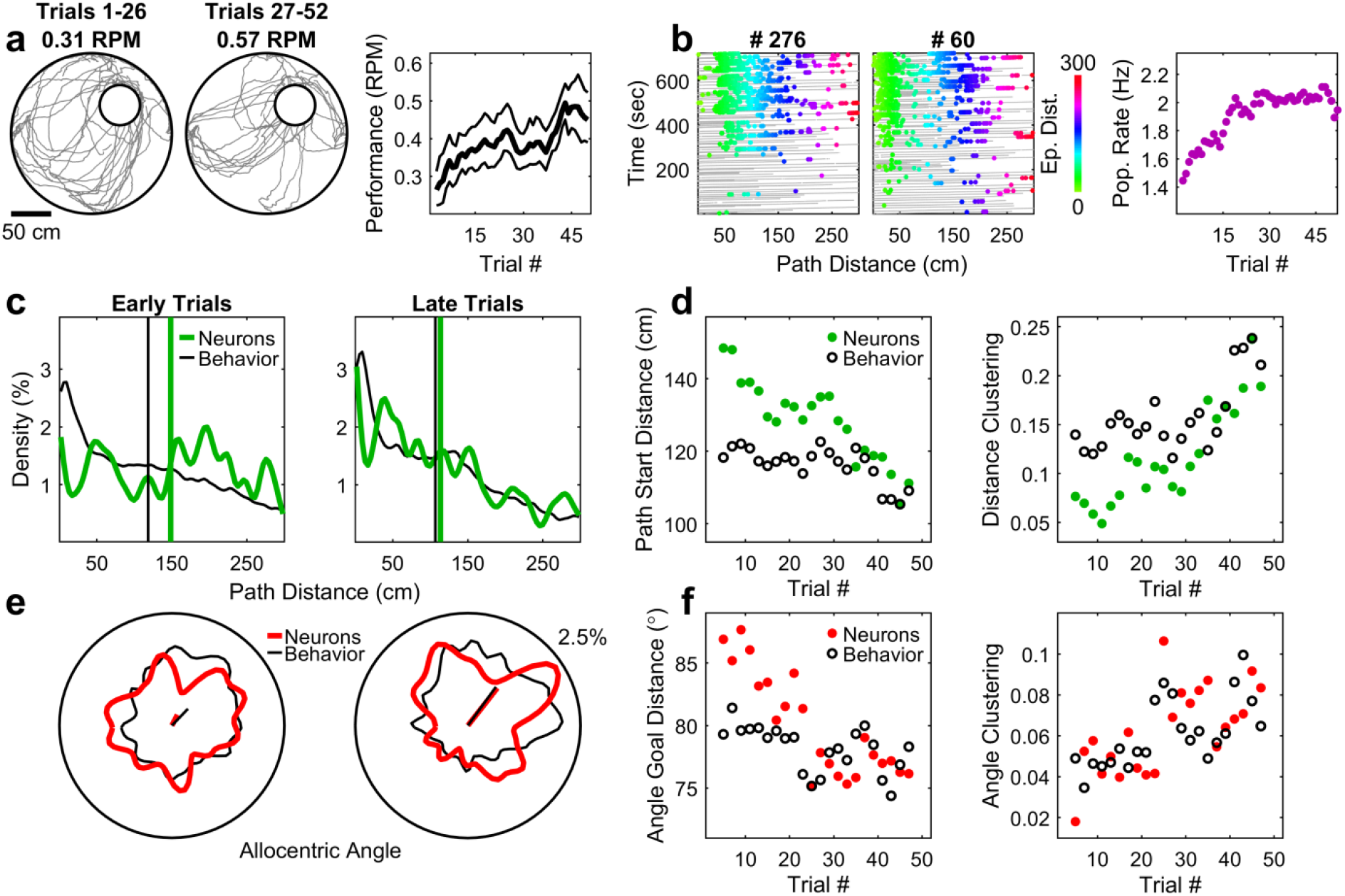
Increased neural clustering correlated with improved behavior within a session. **a,** Paths from trials 1-26 of a session (left) were less efficient than paths from trials 31-52 of the same session (middle). Right, across all sessions, performance increased with increasing trial number (Effect of trial number on Performance: p=1.9×10^-4^, 34 sessions; one-way repeated-measures ANOVA). Thick line: mean; thin lines: 95% confidence interval of the mean. **b,** Left, two example cells which had minimal firing in early trials of a session but increased their firing rates later in the session. Right, the firing rate of active cells increased with trial number (Effect of trial number on Population Rate: p=1.2×10^-3^, 384 units; one-way repeated-measures ANOVA). **c,** Left, distribution of peaks of distance rate maps (green) and occupancy distribution (black) from an early trial (trial 5) across all rats, showing dispersed, fairly uniform distributions (quantified in **d**, right) with median distances of 148 and 118 cm, respectively. Right, distributions of distance peaks and occupancy from a later trial (trial 43) across all rats, showing clustering near the beginning of trials, with median distances of 114 and 107 cm, respectively. **d,** The center of the distribution of tuning curve peaks and behavior both moved closer to the trial beginning with increasing trial number (Neurons: R=-0.91, p=2.8×10^-9^, two-sided *t* test, n=22 trial blocks (every other trial from trial 5 to 47) here and for all other tests in this figure; Behavior: R=-0.69, p=3.9×10^-4^), and the sparsity of these distributions increased with trial number (Neurons: R=0.89, p=2.6×10^-8^; Behavior: R=0.89, p=4.4×10^-8^). **e,** Left, distribution of peaks of angle rate maps (red) and occupancy distribution (black) in early trials, showing dispersed, fairly uniform distributions (quantified in **f**, right), with mean vectors of 63° (mean vector length, MVL, of 0.05) for neurons and 46° (MVL=0.15) for behavior. Right, distribution of angular tuning curve peaks and occupancy in later trials, clustering towards the north-east, with mean vectors of 52° (MVL=0.18) for neurons and 54° (MVL=0.22) for behavior. **f,** The angle goal distance (see Methods) for angular tuning curve peaks and behavior decreased with increasing trial number (Neurons: R=-0.86, p=3.4×10^-7^; Behavior: R=-0.55, p=7.6×10^-3^); the sparsity of these distributions increased with trial number (Neurons: R=0.67, p=7.1×10^-4^; Behavior: R=0.65, p=1.0×10^-3^).

Next, we investigated changes in the distance fields as a function of experience. To isolate the effect of experience on distance tuning, we analyzed 88 cells with significant tuning for path distance, but not allocentric angle (Extended Data Fig. 21). We computed the cross-correlation between the GLM-derived path distance maps from the first and second halves of a session (see Methods). The distribution of the peak lag of these cross-correlations was significantly negative (median of −7.5 cm) indicating of a net backward shift comparable to that seen on liner paths in the real world ^13–18^, though a small proportion of cells did shift forward with experience.

The above results showed rapid experiential plasticity of individual neuron’s receptive fields (see Methods). We then asked if the neural ensemble responses, especially their clustering (Fig 2e-g), also showed any experiential dynamics. Compared to early trials, in the later trials the allocentric spatial rate map peaks clustered towards the unmarked reward zone (Extended Data Fig. 22). The path distance peaks also clustered with experience, but towards the beginning of trials (Fig. 4c, d), rather than the reward zone. Finally, the angular peaks became more clustered with experience, towards the north-east quadrant where the hidden reward zone was located (Fig. 4e-f). These changes in receptive field clustering were accompanied by similar shifts in behavioral biases within the corresponding variables. For all three variables, the measure of central tendency and the sparsity of the distribution showed nearly identical changes with experience – qualitative and quantitative – between neural responses and behavioral measures (Fig. 4c-f).

Qualitatively similar effects were present within data from each rat (Extended Data Fig. 24). Thus, neural ensemble responses increased their overrepresentation of navigationally relevant variables with experience. While distance coding tended to cluster at start locations, spatial coding clustered near the reward site, and angle coding clustered towards the reward quadrant. Specifically, all clustering was near the corresponding regions of high occupancy, and not necessarily near regions of reward, consistent with theories of associative Hebbian plasticity^32^.

To assess the temporal relation between changes in neural coding and changes in behavior we computed the cross-correlation between the neural clustering and behavioral clustering measures as a function of performance. Data was split into two groups containing sessions with high or low behavioral performance (Extended Data Fig. 23). While the neural correlation with behavior was highest in high-performing sessions, neural responses tended to precede behavior in low-performing sessions. This may indicate a differing relationship between hippocampal neural responses and behavior at different stages of learning: a weak and delayed relationship when performance is low, allowing neural responses to drive behavior; and a strong and rapid relationship when performance is high, allowing behavior to be encoded in neural networks.

Finally, we evaluated tuning specificity and its experience-dependence using population vector decoding (Fig. 5, see Methods). For path distance, decoding accuracy was high even in early trials at small distances (Fig. 5b, c), where the highest density of distance fields were centered. In early trials, error increased dramatically after 150 cm, which approximately corresponds to the distance between the goal and the farthest start position. The decoding error for path distance and allocentric angle decreased substantially within the first ~10 trials, particularly at large distances where there were fewer distance fields. Notably, with experience, even large distances had relatively small decoding error, less than expected from simple accumulation of error with distance via path integration. We also computed the population vector overlap between rate maps generated from all the cells in the early trials compared to later trials. This showed significant consistent backwards shift (Fig. 5c, see Methods), demonstrating that distance coding on a population level became more anticipatory with experience. Similarly, for angle, decoding near 45 degrees was near asymptotic levels in early trials with the improvement coming from angles that were less represented in the neural population (Fig. 5d-f). Angle decoding at all directions also dramatically improved with experience, but with subtle and varied anticipatory shifts (Fig. 5f, right), likely owing to the variability of turning behavior within and across sessions. .

**Fig. 5.**
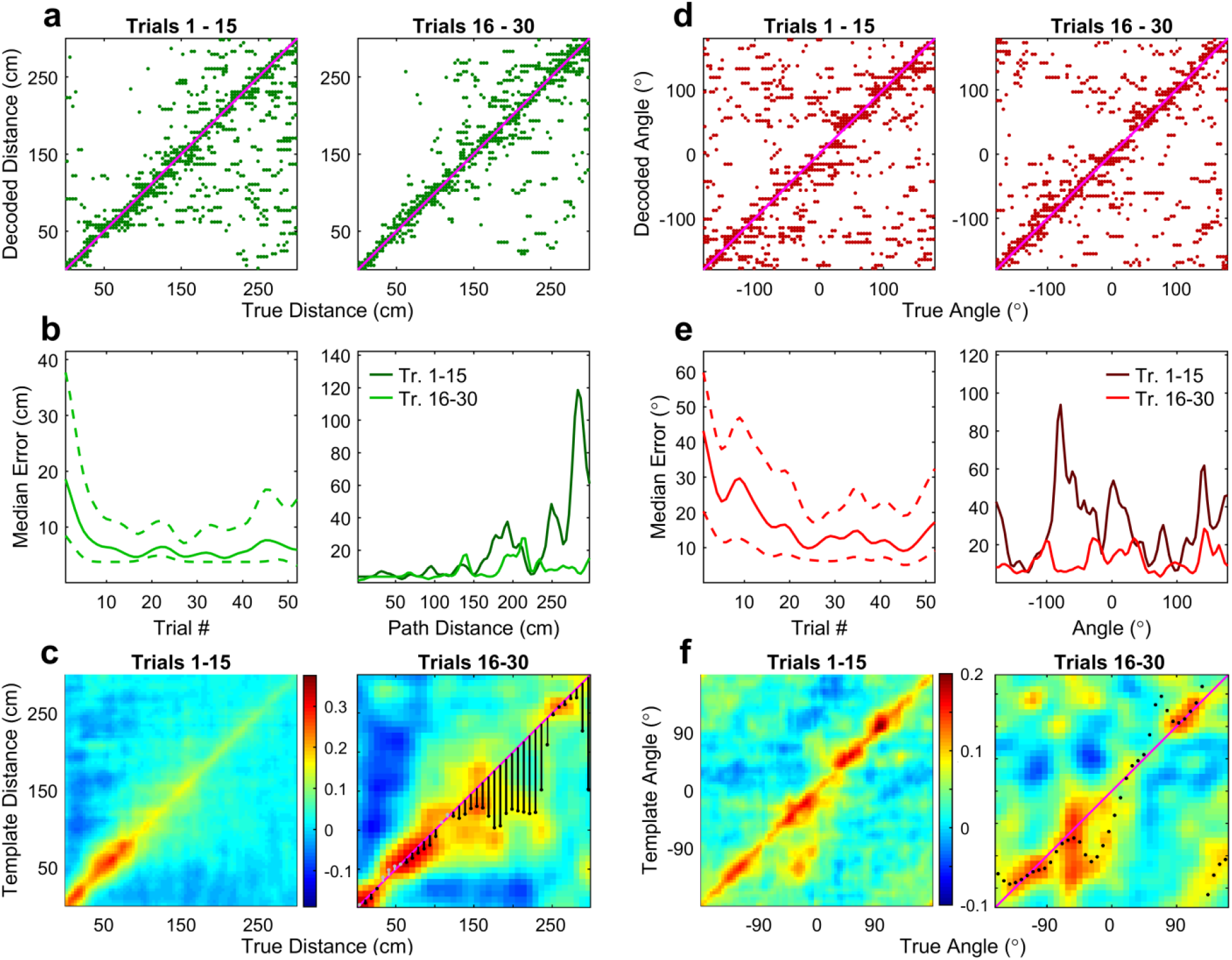
Population vector decoding of path distance and allocentric angle improves with experience. **a,** Decoded distance versus true distance for trials 1-15 (left) and trials 16-30 (right). **b,** Left, median absolute decoding error across all distances as a function of trial number, demonstrating dramatic improvement within the first few trials (Effect of trial number on median error: p=0.01, 80 distance bins, one-way repeated measures ANOVA; Error on trials 1-15 compared to error on trials 16-30: p=1.1×10^-10^, n=1200 (trial, bin) predictions, Wilcoxon rank-sum test). Dotted lines indicate the 95% confidence interval of the median (n=80 distance bins). Right, median absolute decoding error as a function of path distance for trials 1-15 (dark green) and trials 16-30 (light green). Decoding error is low at small distances even in early trials. Most improvement comes at larger distances (Interaction effect between distance bin and trial group: p=3.1×10^-11^, two-way ANOVA, see Methods). **c,** Path distance population vector overlap (see Methods) between entire session activity and activity in trials 1-15 (left) or between trials 1-15 and trials 16-30 (right). Lines and dots mark the smoothed peak correlation on the right-hand plot. Black lines indicate a predictive shift and gray lines indicate a postdictive shift, with a mean value of 15 cm. **d,** Same as **a** but for allocentric angle. **e,** Same as **b** but for allocentric angle. Left, decoding error decreases as a function of trial number (Effect of trial number on median error: p=1.7×10^-7^, 80 angle bins, one-way repeated measures ANOVA; Error on trials 1-15 compared to error on trials 16-30: p=5.6×10^-14^, n=1200 (trial, bin) predictions, Wilcoxon rank-sum test), but not as dramatically as path distance. Right, Decoding error near 45 degrees is smallest, even in earlier trials (Interaction effect between angle bin and trial group: p=0.02). **f,** Same as **c** but for allocentric angle. The best decoded angles span 0 to 90 degrees. Right, experiential shift in angle representation was fairly modest and varied as a function of angle (mean shift −3.2 degrees), which could be due to different turning behavior at specific angles or different turning biases across sessions.

## Summary and discussion

The goal of this study was to determine the nature of hippocampal responses and their potential contribution to navigation, in purely visually guided navigation task, where all other cues, including olfactory and vestibular cues were uninformative. This is similar to the vast majority of primate and human neurophysiology studies of hippocampal function where only visual cues are spatially informative^20,45–47^. Thus, these experiments in rodents help bridge the gap between rodent and human studies, and reveal several similarities^17,48^.

During random foraging-like tasks in VR, hippocampal spatial selectivity is quite weak in primates^45^ and humans^48^, instead showing schema-like responses^20^. Analogously, we find that hippocampal representations of space in a purely visually guided two-dimensional virtual navigation task are markedly different, and more complex, than those observed in previous studies of hippocampal place cells in the real world, where multiple sensory modalities contribute. Allocentric spatial selectivity is far less than that seen during RW foraging tasks in rodents, and similar to that during random foraging in the same VR^25^. Thus, navigational task demand did not have a large impact on allocentric spatial selectivity. Our results would be crucial for understanding the contribution of other sensory cues to hippocampal responses and behavior in a variety of virtual and real world conditions^49,50^.

Our demonstration that rats can execute the virtual navigation task without robust place cells is consistent with the finding that mice lacking robust allocentric place cells^51,52^ can navigate reliably. While our experiments show navigational memory trace in CA1, the upstream areas^53^, e.g. CA3^54^ or the entorhinal cortex^55^ could also contribute. Consistent with computational models of NMDAR-mediated plasticity in areas beyond hippocampus^17,56^, enhancement of NMDAR-mediated plasticity generates larger place fields^57^ and grid cells^58^ greater experiential field plasticity of grid cells^59^ and greater navigational performance^60^.

In contrast to the weak allocentric spatial selectivity, we found very strong allocentric angle and path distance tuning in hippocampal cells. This could be because, in our task, the only spatially-relevant cues available to the rat are visual information and podokinetic (step-counting) cues, and these are fairly dissociated due to the multiple start positions. This could facilitate the dominance of path distance and angular selectivity, as opposed to allocentric selectivity, which may require the repeated pairing of these cues with other stimuli, such as olfactory or tactile^25^. Our experimental paradigm thus allows us to dissociate the contributions of distinct sensory-motor mechanisms that support spatial navigation. This does not imply that allocentric spatial selectivity is never used for navigation, but rather that it is not strictly necessary.

Remarkably, the same cell could multiplex and represent all three variables: allocentric space and angle, and path distance. This supports the hypothesis that the hippocampus is required for learning a schema, that includes both the spatial map and non-spatial components, for accurate navigation^61^. Early studies indicated that hippocampal representations in two dimensions were scalar in nature, dependent only on position of the rat^62^. Subsequent studies showed that the majority of place cells are directionally selective in one dimension, and a significant fraction of place cells encoded allocentric position and head direction in two dimensions^37,63–65^, another example of multiplexed spatial representation. The first example of simultaneous representation of distance and direction was found in virtual reality, when neurons encoded distance in two^23^ or three directions of motion^25^. Similar encoding of distance and direction was found in the real world tasks requiring navigation^26,66^. Our findings are consistent with these and further show that the same neurons encode all three variables. In our experiments, distance coding independent of allocentric position could be interpreted as egocentric, since path distance is purely defined by the beginning of a trial.

## Clustering of neural responses

Clustering of place fields has previously been observed on real world linear tracks at both the beginning and end of the track^23,38^, or primarily near goal locations in other real-world tasks^26,39,40^. However, we see a large clustering of distance fields exclusively at the beginning of the paths. The increased density of firing at short distances may reflect the importance of processing sensory information at the beginning of trials, where uncertainty about the position of the goal is maximal and initial navigational decisions are made.

Unlike random foraging in RW^37,63^ and VR^37^, where the peaks of angular fields were uniformly distributed, the distribution of angular field peaks was clustered towards the hidden reward zone. This indicates that hippocampal angular tuning can be influenced by an un-cued, remembered location^26^, even in the presence of salient visual cues. The goal-centered clustering of angular tuning is complementary to the start-centered clustering of distance tuning, but associative Hebbian plasticity could underlie both phenomena^32^. In both cases, receptive fields clustered towards regions of high behavioral occupancy: near the start for path distance, and toward the goal for allocentric angle. These different reference points are consistent with navigational strategies in the wild which rely on distance from the start position and the direction of the goal zone^26^. Indeed, navigational performance was positively correlated with the degree of spatial, distance, and angular tuning, suggesting that these parameters play a role in navigation^35^.

## Models of synaptic plasticity and receptive field plasticity

The clustering of distance fields at the beginning of trials is also supported by biophysical models of navigational learning ^12–15,17,30–32^ via Hebbian, NMDAR-dependent, temporally asymmetric, spike timing-dependent plasticity (STDP)^30–32^, which predict that with experience, place fields should increase their firing rates and become more anticipatory, i.e. shift backwards towards the start position. These predictions have been extensively supported during linear track running in the real world^13–18^, but are not tested in two dimensions or during a navigational task. Since we are studying well trained rats which had performed several sessions in the same environment, shift of place fields towards the start position would accumulate across days, resulting in a large clustering of place fields at the start position, precisely as observed here. Similar clustering has been observed in one-dimensional tracks in the real world, but this occurs at both the beginning and end of the track^38^. This could be due to the presence of olfactory or reward-related cues that may fix place fields in allocentric space^23,25^. This is also the first evidence for task-related clustering of path distance and allocentric directional tuning, consistent with models of learning^32^ via the associative component of NMDAR-dependent plasticity^28,29^.

Models of STDP also predict a change in the shape of the receptive fields, making them negatively skewed^13,17,67^, which has been observed in subthreshold membrane potential of place cells ^68,69^. However, the asymmetry is the third moment of rate-distribution and hence very sensitive to the detection of the field edges. The episodic distance fields are often multipeaked and rarely reach below 20% of the peak firing rate^13^, which thus precludes the computation of their asymmetry or experiential plasticity. Further, models of place field plasticity show that spike frequency adaptation results in smaller asymmetry of the extracellular spikes compared to subthreshold membrane potential^13,17,67^and seen experimentally^68,69^. Hence, changes in firing rate and center of mass provide more robust extracellular measures of Hebbian plasticity with experience. The backwards shift in distance fields observed here is larger compared to that seen on linear tracks in the real world ^13–18^. This could arise due to differences in task demand in our experiments, and the presence of proximal cues in the RW that anchor neural responses. On the other hand, the time course of experiential plasticity here is slower than that in one dimension ^13,14,17^. This is to be expected because, in one dimensional tasks, the rat experiences exactly the same path in every trial, whereas here rats start from four (or eight) different start positions, so each start position is experienced once every four (or eight) trials. Further, on one dimensional paths place fields only encoded position, whereas here neurons encode three different variables, and each of them show different forms of experiential plasticity. Finally, there is greater trial to trial fluctuation in behavior in our experiments than on narrow linear paths. These mechanisms could slow down the time course of anticipatory shift.

These experiential changes in neural responses occurred every day, even in well trained rats, suggestive of reconsolidation^70^. This is similar to the experiential changes in neural responses not only in the novel environments ^15,18^ but also familiar environments ^13,14,18^ and the magnitude of changes were comparable. This is consistent with computational hypotheses ^17,71^ and experimental findings suggesting ^72,73^ that a certain portion of the hippocampal memory trace is pruned during the slow wave sleep after the experience, resulting in improved signal to noise ratio of memories, and improved performance the next day.

Notably, with experience, there was also a significant increase in the number of cells that were active in the maze^14^. This cannot be explained directly by STDP, which requires spiking. The activation of CA1 may have been inherited by STDP related changes in the presynaptic structure such as CA3^16,74^ or entorhinal cortex^75^, though this is difficult^53^. Alternatively, this plasticity could arise due to dendritic spike^76^ or dendritic plateau potential^77^ dependent, NMDAR mediated plasticity within CA1^17,78,79^.

Improved receptive field selectivity would improve behavior. Conversely, better behavior, i.e. more direct, systematic paths would result in greater receptive field plasticity. This would predict that the two should be correlated with receptive field improvements showing a small lead over behavioral improvement. This prediction was strongly supported by the data – trial by trial fluctuations between performance and neural firing rates, as well as various measures of neural selectivity were strongly and significantly correlated, with neural responses preceding behavior in many cases, indicative of a causal relationship. Interestingly, this causal effect had the largest magnitude in low-performing sessions, while the correlation with behavior was highest in high-performing sessions across multiple measures. This may indicate a differing relationship between hippocampal neural responses and behavior at different stages of learning.

## Relationship to path integration and generalization

Are the distance and direction selective responses related to episodic memory, path integration, or some other abstraction? Place cells show allocentric place selectivity, not distance selectivity in standard, real world experiments, e.g. linear tracks^42^, while distance^23–26,36,66,80^ and time^22,23,41,81^ selectivity has been reported in specific conditions.

In a previous study, distance selectivity appeared when the start box was moved to different locations between trials^36^. This was restricted to a short distance from the box or the rewarded position, and did not generalize to both movement directions. This could arise by some form of path integration that does not manifest in standard tasks, or by olfactory and tactile stimuli of the box. Robust distance coding has been observed in the star maze in the real world^66^, similar to the coding observed in VR when spatially informative olfactory and tactile cues were eliminated^25,42^. This could arise by path integration through a specific set of podokinetic (foot-step) cues or optic flow, since the vestibular cues are largely eliminated in VR. Indeed, when head fixed mice ran on a treadmill in darkness, place cells encoded the distance covered^24^ further supporting podokinetic integration. Additionally, rats running in a fixed place during a memory task had distance^22^ and time^81^ selective cells, rats running in VR had multiplexed responses encoding distance and direction^23,25^, bats flying towards a rewarded pillar had distance and direction selective responses^26^, and visual cue-dependent distance selectivity was found in head-fixed primates^20^.

Our results are broadly consistent with these findings, with important differences. Unlike many real-world experiments, there were no spatially informative olfactory, tactile, auditory or vestibular cues in our virtual maze, nor any specific cues at the hidden reward zone, either before or during the experiment^26,82^. Further, the visual cues were entirely different from each start position in our experiments, and rats had to behave differently to successfully navigate. Thus, the distance selectivity could not arise due to specific movements, vestibular or nonspecific cues, while overcoming the differences in visual cues, and must be computed de novo from each start position. A likely mechanism is path integration. But, path integration is thought to require robust vestibular cues, which are missing in VR. We hypothesize that the underlying mechanism of distance selectivity is podokinetic integration or step counting. Remarkably, path integration is thought to be error prone with rapid error buildup, requiring visual or other cues to stabilize the responses. Whereas we found very little error buildup despite the absence of vestibular cues, highly variable behavior, and visual cues providing contrasting information from the four start arms. Thus, hippocampal neurons can robustly integrate variable podokinetic information, while ignoring contrasting information. Such encoding of distance independent of allocentric position or head direction could be interpreted as egocentric or episodic in our experiments, since path distance is purely defined by the rat beginning a trial.

We hypothesize^25,83^ that 1-5 second motifs of activity arise from upstream structures to CA1 such as medial entorhinal cortex^84^. This then combines with multisensory inputs in the hippocampus using associative, somatic, and dendritic spike mediated Hebbian synaptic plasticity^17,78,79^ to generate selectivity to abstract quantities such as space, distance and angle, to support diverse mechanisms including path integration^6^ and multisensory association^17,23,25^. Thus, distance coding could be an abstraction or generalization, that factors out visual, turning or other sensory cues that differ across start positions, yet can integrate podokinetic cues to generate an invariant, behaviorally relevant measure. Because visual cues are tightly linked with podokinetic cues in linear tracks, but not in 2-dimensional navigation, this may underlie the differences in allocentric spatial coding we observe in our virtual reality.

Head direction information is also crucial for path integration and it is thought to arise from integration of vestibular cues, but the strong angular selectivity in our data cannot arise from these, since vestibular cues are minimized. The angular tuning could be responses to specific set of distinct visual cues on the walls, as observed during random foraging^37^, supporting cognitive mapping^5^ or spatial-view responses in primates^85^. But, the clustering of preferred direction towards the hidden reward zone would require additional computations that combine visual cues with reward information^10,39^ and statistical regularities in behavior, e.g. pointing more frequently towards the hidden reward zone, to orientation information with respect to an invisible, abstract point. Similar to the experiential dynamics of distance coding error, the biggest improvements in the angle coding error occurred at angles away from the goal direction.

## Episodic memory: what, when, where?

These neural responses could form the basis of flexible, episodic spatial memory^21^. Episodic memory is commonly thought to require information about what, when and where. These streams need not be distinct, e.g. hippocampus contains information about place (where) and time (when). But, it has been difficult to find evidence for all three measures of episodic memory, simultaneously. In our experiments the “where” information could be provided by the allocentric selectivity and angular selectivity that encode the position and orientation of the rat and the invisible reward zone, defined only by specific combinations of distal visual cues. The clustering of allocentric place cells near the hidden reward zone^10,39^ supports this hypothesis, along with the experience-dependent forward movement^82^ and increase in clustering. These are further examples of abstract computations that generate a novel form of spatial mapping by combining visual cues with reward information and behavioral regularities. Merely knowing the location of the rat and the reward zone is not sufficient to navigate; information about the orientation of the rat with respect to the reward zone is also required. Consistently, a large proportion of neurons encoded the rats’ orientation, and their preferred angles clustered around the hidden reward zone. The “when” information could be provided in part by the distance selectivity, triggered, but not determined by, movement. Indeed, distance selective cells were also selective for time elapsed, with highly correlated tuning. The “when” information needs to be distinct across different start location and across different trials from the same location. The combination of distance and orientation selectivity could provide distinct information about distance and direction from the start position. Further, there was a significant temporal relationship between these variables, such that distance selectivity preceded angular selectivity and allocentric spatial selectivity. This is indicative of an episodic representation, consisting of distance, followed by angle and then reward information. These form a trajectory-specific episode, unique for every path or trial, to successfully navigate.

Experience-dependent changes in the distance, orientation and allocentric positional information could provide distinct information across trials. The time scale of these experience-dependent “when” information is consistent with that observed in the real world, one dimensional paths^13–18^, and with the time encoding cells in the lateral entorhinal cortex^27^, upstream to CA1. Finally, a significant portion of neurons multiplexed and encoded all three codes. Thus, not only the ensemble of neurons but these individual neurons could provide information about “what” or the schema of the entire experience.

These results provide evidence for the sequential, episodic arrangement of simultaneously existing allocentric and path-centric information in hippocampal neurons that is correlated with navigational performance. These neural responses are highly dynamic, which can be explained by computational models of navigational learning via both associative and asymmetric components of synaptic plasticity. Thus, the results help bridge the gap in our understanding about the cellular mechanisms of Hebbian plasticity, hippocampal episodic and allocentric responses, and navigational performance.

## Acknowledgments

We thank C. Vuong, B. Popeney, and D. Aharoni for help with initial development of the experimental paradigm and early data collection. This work was supported by grants to M.R.M. from the W.M. Keck foundation, AT&T, NSF-EAGER, and the NIH-U01. Some results presented in this manuscript were presented in abstract form at the annual Society for Neuroscience conference in 2015, 2016, 2017, 2018, and 2019.

## Author contributions

J.D.C, L.A., and M.R.M. designed the experiments. J.D.C and L.A. performed the experiments. J.J.M. performed analyses with input from M.R.M. J.J.M and M.R.M. wrote the manuscript with input from other authors.

## Methods

### Brief methods

We used a body-fixed virtual reality system in which rats were trained to run to a hidden reward location in the virtual space as described previously^33^. Single units were recorded from bilateral CA1 in well-trained rats (n=4)^25^. Units were manually clustered off-line, and well-separated pyramidal units with a mean firing rate during movement greater than 0.5 Hz were included in all analyses. A total of 384 units meeting these criteria were recorded across 34 behavioral sessions. No systematic differences were observed between data from different rats, so all units were pooled together for analysis. Additionally, no systematic differences were observed between data from sessions with 4 start positions or 8 start positions, so these data were pooled together as well, except for direct comparisons between the two conditions. Selectivity maps for allocentric position, path distance, and head angle were simultaneously estimated using a generalized linear model (GLM) framework^37^. The degree of selectivity was quantified by sparsity, and statistical significance for each unit was assessed using its own shuffled data.

### Subjects

4 adult male Long-Evans rats were implanted with bilateral hyperdrives each containing up to 12 tetrodes per hemisphere. Rats were food and water restricted to motivate performance.

### Behavioral task

Animals navigated in a virtual space using a body-fixed virtual reality system previously described^23,33^. The virtual table was circular and 100 cm in radius, placed in the center of a room measuring 400 x 400 cm. Each wall had a unique visual design on it to provide a rich visual environment (Fig. 1a). The table had a finely-textured pattern to give optic flow without providing spatial information. The table was placed 100 cm above a floor with a black and white grid pattern so rats were able to visually detect and turn away from the edge of the table^33^.

Trials began with the rat in one of 4 (or 8) start positions, at a distance of 5 cm from the table edge facing radially outwards. These start positions corresponded to those directly facing the walls (defined as North, East, South, and West) for sessions with 4 start positions, and angles in the middle of these for sessions with 8 start positions. The hidden reward zone (radius of 20-30 cm) was always located in the North-East quadrant with its center at coordinates *(35.3, 35.3)*.

Rats freely moved about the virtual space until they entered the reward zone. Upon entry, the reward zone turned white, and pulses of sugar water were delivered at 500 ms intervals, accompanied by auditory tones for each pulse. This continued until 5 rewards were delivered or the rat exited the reward zone, ending the trial. At trial end, the visual scene was turned off and a blackout period of 2-5 seconds ensued. Rats were teleported to a new randomly chosen start position during this period. The visual scene was then restored, and a new trial began.

To encourage the use of a navigational strategy rather than a stereotyped motor response, experimental conditions were occasionally changed to define “session blocks” unique to each rat. Specifically, the number of start positions, reward zone size, or the set of wall cues was changed every 2-4 days, yielding a total of 12 session blocks.

The primary measure used to quantify performance was rewards/meter (RPM), equivalent to the inverse of the mean path length. To quantify the improvement in behavior across sessions, we calculated the percent change in RPM relative to the first day in a session block (Fig. 1b, right). To test for statistical significance of the change in performance, the difference between the percent change on consecutive days within a session block was computed (27 such differences), and was subjected to a Wilcoxon sign-rank test. This is a conservative estimate as we include data from up to 4 days in the same condition, while the biggest improvement tended to occur between days 1 and 2 within a session block.

Position in the virtual environment was sampled at a rate of 55 Hz. Throughout the paper, “angle” refers to the viewing angle of the virtual avatar. Angle is purely defined by visual cues, as the rat is body fixed to point in the same direction in the real world frame of reference at all times.

### Electrophysiology

Neural activity was recorded extracellularly from dorsal CA1 using tetrodes. Tetrodes were made from a Nickel-Chromium alloy and insulated with polyimide. Action potentials were detected as described previously and manually sorted into putative neurons, or units^23,25^. Only putative pyramidal neurons were used for analysis, identified by having a high complex spike index (>15) and waveform with a width at half-max at least 0.4 ms. Additionally, only units with a mean firing rate greater than 0.5 Hz during movement (speed>5 cm/s) were included, in order to ensure sufficient data for analyses. Tetrodes were adjusted daily to increase the total number of independent neurons.

### Binning method of computing rate maps

Similar methods as described previously^25^ were used to compute rate maps using the binning method. In all analyses, only behavior and neural activity when the rat was running faster than 5 cm/s outside the reward zone were included. Additionally, only data with a path distance less than 300 cm was included, since longer run distances were rare (<5% of trials).

Spatial rate maps were computed using bins of size 5×5 cm spanning from −100 to 100 cm in both *X* and *Y* coordinates. Occupancy maps and spike count maps were computed, smoothed with a 2-dimensional Gaussian kernel, and then divided to compute the rate. In Fig. 1 and Extended Data Fig. 3, the 2D smoothing kernel had a sigma of 7.5 cm, to directly compare values to previous work. In all other figures, the 2D smoothing kernel had a sigma of 5 cm. Bins with less than 250 ms of occupancy were excluded.

Path distance maps were computed in a similar fashion, with 80 bins of width 3.75 cm spanning 0 to 300 cm, and smoothed with a Gaussian kernel with a sigma of 3.75 cm. Bins with occupancy less than 2 seconds were excluded.

Angular rate maps used bins of size 4.5 degrees from −*π* to *π*, circularly smoothed with a Gaussian kernel with a sigma of 4.5 degrees.

Temporal rate maps (Extended Data Fig. 11) were constructed in a similar fashion. The moment a rat began moving in a trial was defined as time 0. We used 80 bins of width 250 ms spanning 0 to 20 seconds, and smoothed with a Gaussian kernel with a sigma of 250 ms.

Goal distance maps (Extended Data Fig. 12) were constructed in a similar fashion to path distance maps using the same sized bins, but from −300 cm to 0 cm, with 0 cm indicating the moment the rat entered the reward zone.

### Generalized linear model

A generalized linear model (GLM) with logarithmic link function, similar to the one used in previous work^37^, was further developed to estimate the simultaneous contribution of space, distance, and angle to neural firing. For each neuron, spike counts were binned using 100 ms bins. For allocentric space, between 5 and 32 Zernike polynomials were used as basis functions^37^. For path distance, basis functions were the first 10 Chebyshev polynomials of the first kind^86^. For allocentric angle, basis functions were sine and cosine functions with frequencies *n/2π* for *n={1,2,3,4,5}*. These choices of basis functions provided orthonormal sets which spanned the entire parameter range.

Coefficients for these basis functions were fit with the Matlab function *glmnet()*^87^ with an alpha parameter of 0 using 5-fold cross validation and 100 lambda values generated by *glmnet()*. This was repeated for models containing 5 to 32 Zernike polynomials, yielding 2800 possible models for each neuron. The fitness *F* of each model was computed as the mean cross-validated error. To bias model selection towards smooth maps, this fitness was then divided by the coherence of the spatial rate map for each model to get the final fitness *F\**. The coefficients corresponding to the model with the lowest *F\** value were then used to reconstruct the rate maps (see below)

For plotting purposes and calculation of tuning, GLM-derived maps for space, distance, and angle were constructed by evaluating the GLM at the center of the bins described above for binned rate maps. Minimum occupancy values were identical to those for binned rate maps.

Specifically, GLM allocentric space maps R_Space_(X,Y) were calculated as 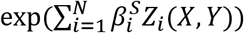, where 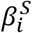 is the coefficient for the *i^th^* spatial component, *Z_i_* is the *i^th^* Zernike basis function, and *(X, Y)* are the centers of the 5 cm x 5 cm spatial bins. Here, *N* is the number of basis functions selected by the above cross-validation process.

Path distance maps R_Distance_(D) were calculated as 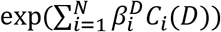, where 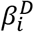 is the coefficient for the *i^th^* distance component, *C_i_* is the *i^th^* Chebyshev basis function, and *D* represents the centers of 3.75 cm-wide distance bins from 0 to 300 cm. Here, *N=10*.

Allocentric angle maps R_Angle_(A) were calculated as 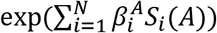, where 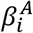 is the coefficient for the *i^th^* angle component, *S_i_* is the *i^th^* sinusoidal function, and *A* represents the centers of 4.5 degree-wide angle bins from −π to π. Here, *N=10*.

The predicted firing rate of a neuron from these GLM-derived maps is the product of all 3 maps with a constant scaling factor: R(t) = c*R_Space_(X(t),Y(t))*R_Distance_(D(t))*R_Angle_(A(t)). Each individual map can be considered akin to an individual “risk factor” for firing. The spatial, distance, and angle curves along with the constant term must be considered simultaneously to derive a predicted firing rate at any time. Consequently, the maximum rate for any individual GLM-derived curve is arbitrary. We therefore present these maps in terms of normalized rates. As sparsity is a scale-invariant measure (see below), the sparsity of normalized rate maps is identical to the sparsity of non-normalized rate maps. When individual GLM-derived maps are presented (Figs. 2, 3, Extended Data Figs. 4, 5, 8, 9, 14, 15), the mean firing rate (*μ*) of the unit is noted, to give an idea of the general activity rate of the unit.

### Quantifying tuning

We used sparsity to quantify the degree of tuning of each rate map. Maps were divided into *N* bins as defined above, and sparsity was computed as

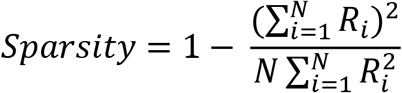

Where *R_i_* is the value of the rate map in the *i^th^* bin.

### Statistics

For determining the statistical significance of rate maps, each unit served as its own control. For each parameter (space, distance, angle), the relevant behavioral vector was time reversed and shifted by *n*s* seconds, for *n = 1 to 60*, where s is the duration of the session divided by 61. Rate maps were re-estimated for this altered dataset, and the resulting sparsity values formed a null distribution for the shifted parameter. Rate maps were considered statistically significantly tuned if the original sparsity exceeded all 60 control sparsity values, yielding an effective statistical criterion of *p<0.017*. This procedure is non-parametric and does not make any assumptions about the nature of the null distributions. Offsets were generated from a continuous range rather than using randomly-drawn offsets to eliminate the possibility of randomly selecting two offset values that are near each other. This ensures that the shuffled distribution is composed of independent data points.

Statistical comparisons were made using the Wilcoxon rank-sum test for non-matched data and the Wilcoxon signrank test for matched data. Significance of correlation values were assessed on the linear correlation coefficient using a T-test.

Unless otherwise specified, all values are reported as the median and 95% confidence interval of the median, in the form *M [L, U]*, where *M* is the median of the data and *L* and *U* are the lower and upper bounds, respectively, of the 95% confidence interval of the median. Unless otherwise stated, error bars in all figures indicate the 95% confidence interval of the median. Confidence intervals for percentages were obtained using the Matlab function *binofit()*.

### Weighted Correlations and Linear Fits

To robustly estimate the relationship between performance and neural tuning (Fig. 3c, Extended Data Fig. 18), we computed both unweighted and weighted correlation coefficients and linear fits. For weighted calculations, each session was weighted by the number of units in that session. To determine statistical significance, we performed a bootstrapping procedure. We resampled the data 10,000 times with replacement to generate a distribution of weighted correlation coefficients. The p-value was the fraction of resampled datasets with a correlation coefficient R <= 0 for Fig. 3c and Extended Data Fig. 18c, or R >=0 for Extended Data Fig. 18a-b.

### Stability of GLM maps

To demonstrate the robustness of the GLM fitting procedure we performed a stability analysis (Extended Data Fig. 5). Tuning curves for the first and second half of sessions were estimated using the GLM procedure described above using data only from trials 1-26 and 27-52, respectively. Valid bins in these restricted rate maps had to have a minimum occupancy > 50 ms for allocentric space, and > 500 ms for path distance and allocentric angle.

Stability is defined as the correlation coefficient between first-half and second-half maps. Additionally, as stated above, all GLM fitting was performed using 5-fold cross validation to mitigate the effects of overfitting. Cells were classified as “tuned” or “untuned” based on their maps estimated from the entire session, as described above (Statistics). Null distributions in Extended Data Fig. 5 were obtained by computing the correlation coefficients between random first-half and second-half maps by shuffling cell identities once.

### Episodic relationship between distance, angle and position codes

To compute the percentage of cells significantly tuned for space, distance, or angle as a function of the progression of a trial (Fig. 2h, Extended Data Fig. 17), we assigned a center coordinate for each cell as the location of the peak of its path distance field derived from the GLM. Percentage tuned as a function of distance was then computed using the distance bins defined for binned rate maps, and smoothed with a Gaussian kernel with a sigma of 15 cm.

The significance of the ordering of tuning to the three variables was tested using a cross-correlation analysis (Extended Data Fig. 17c). To compute significance while controlling for the non-uniform distribution of peak locations, we performed a resampling procedure. All units, significant or not, were assigned a new peak location by resampling with replacement from the original pool of peak locations; the population of these surrogate cells then went through the same procedure described above to generate a null curves and null cross-correlations. This was repeated 5000 times, and the dotted lines in Extended Data Fig. 17c represent the 95% range at each lag for the control data.

### Analysis of variance (ANOVA) for sparsity between conditions

Sparsity is typically negatively correlated with the logarithm of the number of spikes a neuron fires in a session^37^. Thus, we used a two-way ANOVA in Matlab to compare sparsity between virtual navigation (VNT), random foraging in real world (RW), and random foraging in (VR) (Fig. 1e) or between 4- and 8-start VNT tasks (Extended Data Figs. 3d, 6b, 8d, and 14d). Recording condition (VNT, RW, VR, 4-start, or 8-start) was a categorical predictor, and *log_10_(number of spikes)* was a continuous predictor of the sparsity. The p-values reported in the identified figures are for the main effect of recording condition on sparsity.

### Population Vector Overlap and Population Vector Decoding

Population vector overlap and population vector decoding (Fig. 5) were done using binned rate maps for path distance or angle. The “template” vectors in Fig. 5c, left and 5f, left were constructed from all data in trials 1-52 across all sessions and rats. The “true” vectors in Fig. 5 were computed for each trial. The correlation coefficient between each pair of distances (or angles) was then computed for each trial. The matrices plotted in Fig. 5c, left and 5f, left were constructed by averaging the trial-wise matrices for trials 1-15 (left).

To construct the population vector overlap matrices in Fig. 5c, right and 5f, right, population rate maps were constructed using bins twice the width as defined for binned rate maps, to compensate for the smaller amount of data in single trials (7.5 cm for distance, 9 degrees for angle; minimum occupancy of 200 ms in a given trial). “Template” rate maps were constructed from all data in trials 16-30, and “True” rate maps were constructed from all data in trials 1-15.

Population vector decoding (Fig. 5) was done for each bin by picking the distance (or angle) corresponding to the highest correlation in that bin. In Fig. 5c, right and 5f, right, the decoded value was then smoothed with a Gaussian kernel with a sigma of 1 bin (7.5 cm for distance, 9 degrees for angle).

### ANOVA for decoding accuracy across trials and bins

We used a two-way ANOVA in Matlab to compare decoding accuracy in different distance or angle bins across different trials (Fig 5b, right and 5e, right, respectively). Trial number was set to have random effects, and bin number was a continuous predictor of decoding accuracy.

### Average Paths

To construct the average path from each start position to the goal location (Extended Data Fig. 1a), individual trials were grouped according to start position and then interpolated as follows.

For a given start position, define *D_max_* as the distance traveled in the longest path (in cm). For each trial originating from that start position, define *P* as the *X* and *Y* coordinates of the path, and *D_1_* as the cumulative distance traveled in that trial only, normalized to have a maximum value of 1. *D_2_* is defined as a linearly spaced vector from 0 to 1 with *D_max_* datapoints. *P_interp_* is then computed using the Matlab function *interp1(D_1_, P, D_2_)*. The average path is then computed as the median of all interpolated paths.

### Path Correlation

To estimate across-position path correlations (Extended Data Fig. 1b, 18), we computed correlation coefficients of rotated occupancy maps for each start position. Rotated occupancy maps were computed by rotating all paths originating from a given start position such that the initial heading was −90° (south), and then discarding the first 3 position bins (to reduce spurious correlation from low speed at the beginning of trials). Occupancy in 10×10 cm bins was then computed without smoothing to make the rotated occupancy map for each start position. The larger bin size was used to compensate for the reduction in the amount of data resulting from restricting the analysis to individual start positions. The “across-position path correlation” was then the mean correlation coefficient across all pairs of maps from different start positions.

Within-position path correlations were calculated in a similar manner as across-position correlations. First the rotated occupancy map for a start position was computed. Then we repeated the process by resampling individual trials with replacement to construct a new occupancy map. This was repeated 100 times. The “within-position path correlation” was then defined as the 1^st^ percentile (lowest) correlation coefficient between the original occupancy map and the resampled maps.

### Occupancy index and speed Index

To quantify behavioral performance, we calculated the occupancy index and speed index (Extended Data Fig. 2g, h) as follows. Position was binned using 5×5 cm bins. For each session, the occupancy as a function of radial distance from the reward was calculated as the mean occupancy time of bins falling within 6 cm radial bins. Speed as a function of radial distance was computed in a similar manner.

To compute occupancy and speed indices, null behavioral data was generated for each session. For sessions with 4 start positions, the path for each trial was rotated by 90, 180, and 270 degrees. For sessions with 8 start positions, each path was rotated by 45, 90, 135, 180, 225, 270, and 315 degrees. Rotated paths were truncated after first crossing into the reward zone, if applicable. Null radial occupancy and speed were then computed from these rotated data sets. Occupancy index was then defined as:

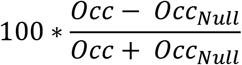

Where *Occ* is the original occupancy distribution and *Occ_Null_* is the null occupancy distribution.

Speed index was defined in a similar way.

### Decomposing maps, goodness of fit, peak index, peak width

To decompose path distance maps into individual distance fields (Extended Data Fig. 9), we iteratively fitted mixtures of Gaussians to the rate maps. First, the rate map was smoothed with a Gaussian kernel with a sigma of 7.5 cm (2 bins). Then a mixture of *N* Gaussians with a constant offset was fitted to the curve where *N* is defined as the number of distinct peaks above 25% of the maximum value. Distinct peaks were defined as peaks with a value below the 25% threshold between them. The reconstructed curve was then re-estimated using the same procedure (without smoothing), and this was repeated until the number of components did not change. The results were robust to small changes in the specific values of the above parameters.

The procedure for decomposing allocentric angular rate maps (Extended Data Fig. 15) was similar, except mixtures of Von Mises functions^88^ were used, and the original curve was circularly smoothed with a sigma of 13.5 degrees (3 bins).

Goodness of fit (Extended Data Figs. 9, 15) was calculated as the correlation coefficient between the original rate map and the map reconstructed from its fit components.

Peak index (Extended Data Figs. 9, 15) was computed for each distance peak, and calculated as the ratio of *A/C*, where *A* is the amplitude of the fitted component and *C* is the constant offset of the fitted curve.

For path distance, peak width (Extended Data Fig. 9) was defined as the sigma value for each fitted component.

For allocentric angle, peak width (Extended Data Fig. 15) was defined as the width of each fitted component at 50% of the component’s amplitude (i.e. full width at half max).

### Inclusion Criteria for Different Path Lengths

To test if the aggregation of distance field peaks was the result of shorter distances being oversampled, we compared the distribution of path distance field peaks for trials of different lengths (Extended Data Fig. 10). Data was separated into 4 groups, comprising of data from trials of 0-75 cm, 75-150 cm, 150-225 cm, and 225-300cm. Binned rate maps were computed as described above, but with a larger smoothing kernel (sigma = 7.5 cm) and a minimum occupancy of only 1s, to adjust for the inclusion of less data. Within each distance range, only cells that met occupancy criteria in at least 75% of bins were included.

To compute the distributions of distance peaks in the bottom panels of Extended Data Fig. 10, peaks were binned using 80 equally spaced bins from 0 cm to the maximum distance for that range, and smoothed with a Gaussian smoothing kernel with a sigma of 3 bins. Here, the smoothing window was defined using bins rather than cm, to allow unbiased comparison of these distributions.

### Rate Modulation Index

Rate modulation index (Extended Data Fig. 20d) was computed as 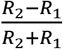, where *R_1_* is the mean firing rate of a cell across trials 1-26, and *R_2_* is the mean firing rate of a cell across trials 27-52.

### Experience Dependence

For analyses of experience dependence (Fig. 4c-f, Extended Data Figs. 22-24, but not Extended Data Fig. 21, see below), occupancy and spikes were binned for individual trials using bins as defined in “Binning method of computing rate maps” above, and smoothed across trials with a Gaussian kernel with a sigma of 4 trials. The smoothed spike maps were divided by the smoothed occupancy maps to produce trial-by-trial rate maps. Minimum occupancy was 40 ms for path distance and allocentric angle, and 20 ms for allocentric space.

Trial-by-trial rate maps for path distance and allocentric angle were decomposed as above to identify peaks, which were then analyzed for Figure 4c-f.

Allocentric space peaks in Extended Data Fig. 22 were defined as local maxima greater than 20% of the peak rate.

Peak density plots in Figure 4c, 4e, and Extended Data Fig. 22a were constructed by binning peaks using bins as defined in “Binning method of computing rate maps” above for each trial. Values in each bin in this distribution were then divided by the total number of cells with defined rates in that bin. This normalization ensures that experience-dependent changes in the distribution of rate map peaks are not artificially driven by experience-dependent changes in the distribution of occupancy. Finally, the distribution for each trial was normalized to sum to 100%.

Single unit shifts in distance tuning curves (Extended Data Fig. 21) were evaluated on GLM-derived rate maps for trials 1-26 and 27-52, as in Extended Data Fig. 5. To isolate the effect of experience on distance tuning only, we selected cells which were i) significantly tuned for distance; ii) not significantly tuned for angle; and iii) included sufficient spiking data in both halves to construct rate maps. 91 cells met criteria (i) and (ii), and 88 cells met all three criteria.

### Experience-dependent clustering analyses

To quantify the experience dependent changes in both neural responses and behavior for allocentric space, we computed two measures: distance from reward and spatial clustering. To compute distance from reward, the distributions shown in Extended Data Fig. 22 were reparametrized by computing the radial distance between any position and the reward zone (Extended Data Fig. 22a, top). This measure was binned using 40 bins of 3 cm width from 0 to 120 cm to generate a distribution of radial distance. Distance from reward was defined as the center of mass of this distribution (Extended Data Fig. 22b, left). Spatial clustering was defined as the sparsity (defined above) of this distribution (Extended Data Fig. 22b, right). The advantage of these measures is that they can be applied to both behavioral and neural data to allow not just qualitative, but quantitative, comparisons between them. These same measures were applied to the distribution of the peaks of the allocentric spatial rate maps (Extended Data Fig. 22a, bottom) to estimate their distance from reward and clustering.

Similarly, we computed the distribution of path distance for both behavior and peaks of path distance rate maps (Figure 4c), and defined distance from start and distance clustering (Figure 4d) in the same way as above.

For angular tuning, we reparametrized the angle distributions (Figure 4e) as the absolute difference between any angle and 45 degrees (corresponding to the northeast quadrant where the hidden reward zone is located). This angular difference was binned using 40 bins of width 4.5° from 0 to 180° to generate a distribution which was quantified as above to compute angular distance from goal and angular clustering (Figure 4f).

### Temporal relation between neural and behavioral changes

To assess the temporal relation between changes in neural coding and changes in behavior we computed the cross-correlation between the neural clustering measures and the behavioral clustering measures (Extended Data Fig. 23, bottom row). Statistical significance was assessed by a shuffling procedure. The neural and behavioral curves were shuffled with respect to trial number to create control cross-correlations. This was repeated 5000 times to create the 99% range indicated by the dotted lines in Extended Data Fig. 23, bottom row.

---

**Extended Data Fig. 1.**
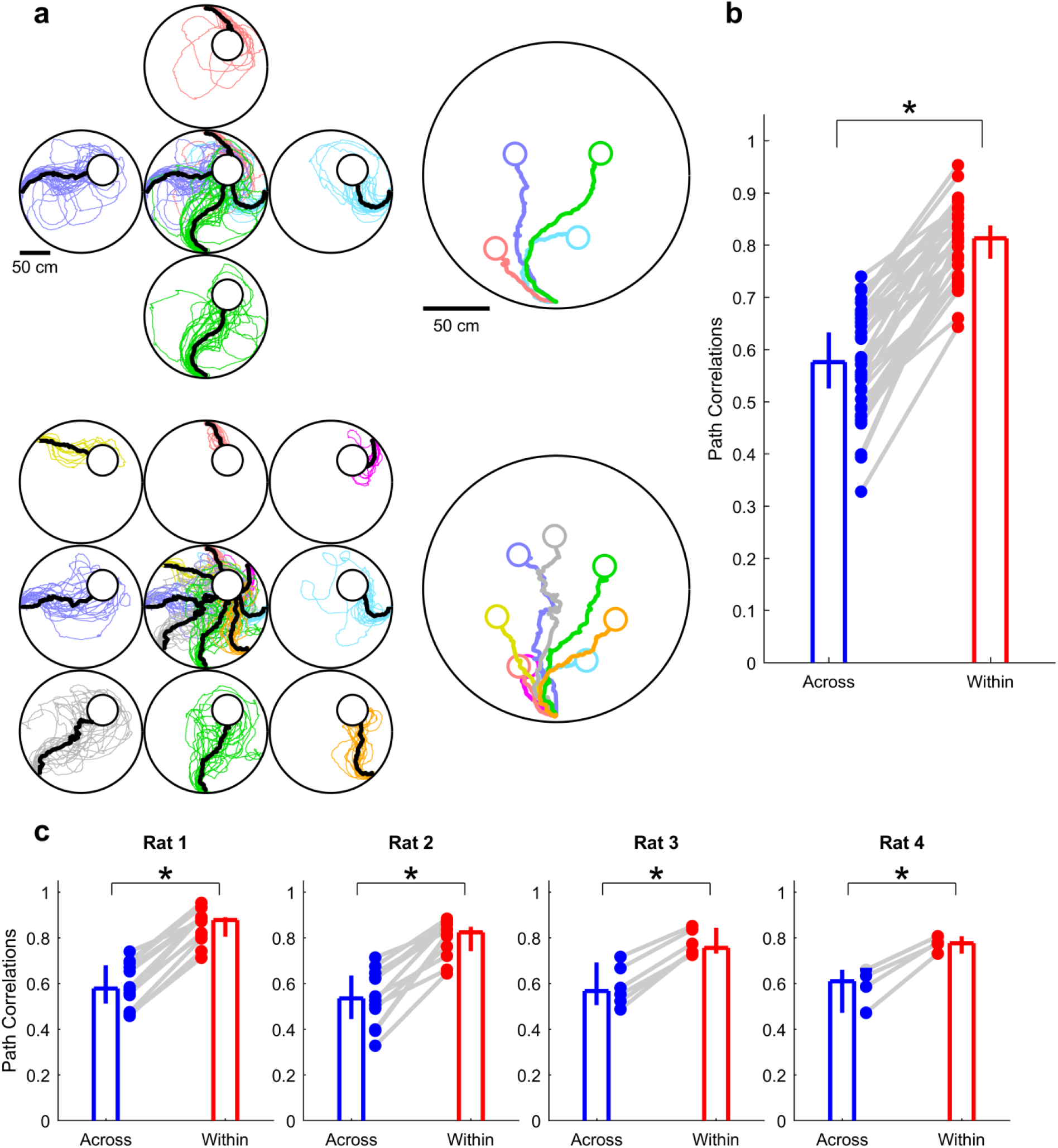
Rats use a place navigation strategy to navigate. **a,** Top, left, individual trials (thin colored lines) and mean path from each start position (thick black lines) for a single behavioral session with 4 start positions. Paths are color coded based on start position. Right, all mean paths rotated to begin at the same point and heading, to illustrate that rats take unique paths from each start position Paths are color coded to match the colors in the left panel. Bottom, same as **a** but for a different behavioral session with 8 start positions. **b,** The path correlation (see Methods) was significantly smaller (p=1.9×10^-7^, One-tailed Wilcoxon sign-rank test) across start positions (0.58, [0.53, 0.63]) compared to within start positions (0.81, [0.77, 0.84]), n=34 sessions for all statistics. **c**, As in **b**, the across start position correlation was smaller than the within start position correlation for each individual rat in the study. Rat 1: Across (0.58, [0.51, 0.68]) vs Within (0.88, [0.80, 0.89]), p=2.4×10^-4^, n=12 sessions). Rat 2: Across (0.53, [0.44, 0.63]) vs Within (0.82, [0.74, 0.85]), p=2.4×10^-4^, n=12 sessions). Rat 3: Across (0.56, [0.50, 0.69]) vs Within (0.76, [0.73, 0.84]), p=1.6×10^-2^, n=6 sessions). Rat 4: Across (0.61, [0.47, 0.66]) vs Within (0.78, [0.73, 0.81]), p=0.06, n=4 sessions). One-tailed Wilcoxon sign-rank test used throughout.

**Extended Data Fig. 2.**
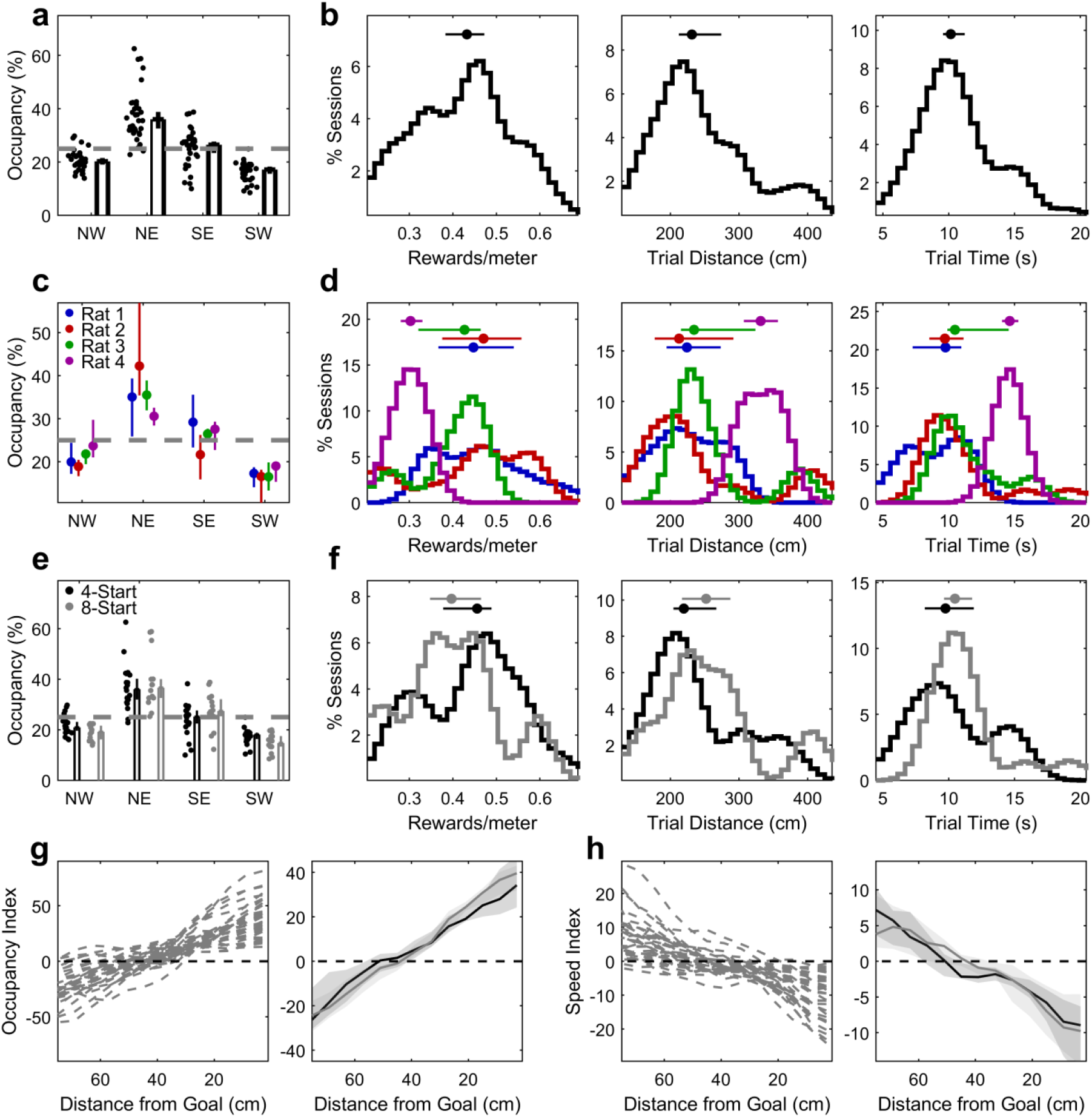
Quantification of behavior as a function of reward location. The hidden reward zone was in the northeast (NE) quadrant. **a,** The percentage of time rats spent in the NE quadrant (33, [31, 37]%) was significantly greater than chance (p=2.8×10^-4^, Wilcoxon sign-rank test), and greater than all other quadrants (NW: 20, [19, 22]%; SE: 28, [26, 30]%; SW: 17, [16, 18]%). **b,** Left: Across all sessions, the median performance in rewards/meter was 0.43, [0.38, 0.47]; Middle: The median trial distance was 230, [210, 260] cm; Right: The median trial time was 10, [9.5, 11] s of movement. **c,** Same as **a** but plotting the median and 95% confidence interval of the median for individual rats. **d**, Same as **b** but for individual rats. For all measures, Rat 4’s statistics (purple) are significantly different from Rat 1 (p=1.1×10^-3^) but not different from Rat 2 (p=0.06) or Rat 3 (p=0.11), using Wilcoxon rank-sum test for all comparisons. **e**, Quadrant occupancy as in **a**, split between 4-start sessions (black) and 8-start sessions (gray), exhibiting similar characteristics. **f**, Behavioral measures from **b**, split between 4-start sessions (black) and 8-start sessions (gray). No significant differences exist between the conditions in any measure. Rewards/meter: 4-start (0.46, [0.34, 0.49]), 8-start (0.40, [0.35, 0.46]), p=0.35. Trial Distance: 4-start (220, [205, 267]), 8-start (252, [217, 297]), p=0.30. Trial Time: 4-start (9.75, [8.47, 12.0]), 8-start (10.5, [9.9, 11.7]), p=0.27. **g,** The quadrant based analysis can be influenced by nonspecific parameters, such as behavioral differences from each start position. Hence, we developed novel analysis based on the distance from the goal, regardless of start position. Left, occupancy index in individual sessions (see Methods) as a function of radial distance from the goal location. Effect of Distance from Goal on Occupancy Index: p=1.3×10^-12^, 34 sessions; one-way repeated-measures ANOVA. Right, population average, showing rats spend more time near the goal than farther from it, compared to chance. The darker line and shading is the median and 95% confidence interval of the median for 4-start sessions, and the lighter line is the same for 8-start sessions. **h,** Left, speed index in individual sessions (see Methods) as a function of radial distance from the goal location. Effect of Distance from Goal on Speed Index: p=2.0×10^-9^, 34 sessions; one-way repeated-measures ANOVA. Right, population average, showing rats run slower near the goal than farther from it, compared to chance. Color conventions are as in **e**. n = 34 sessions for all combined statistics; n=20 sessions for 4-start statistics; n=14 for 8-start statistics.

**Extended Data Fig. 3.**
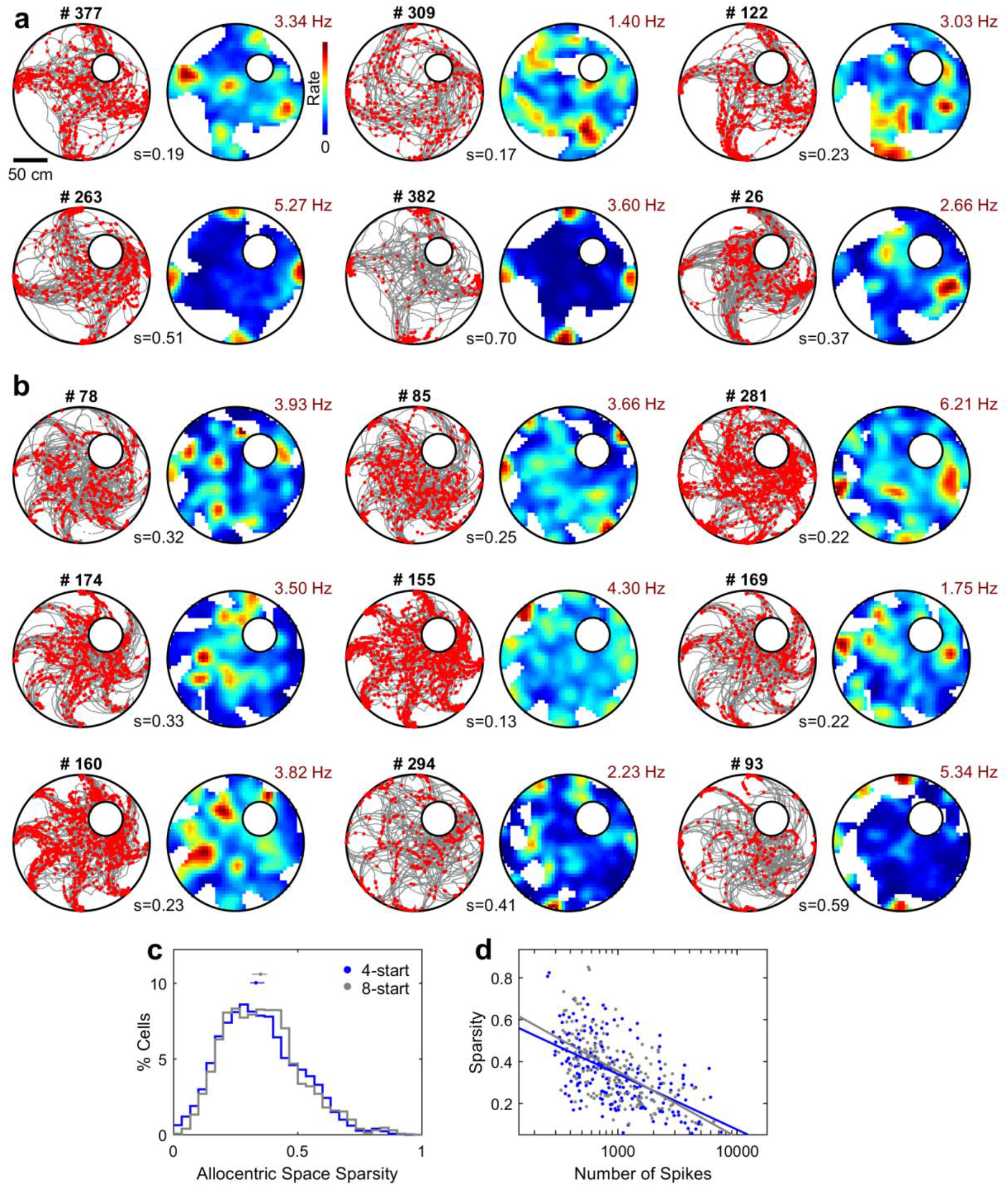
Additional examples of spatial tuning in 4- and 8-start navigation tasks. **a,** Example units as in Fig. 1b-d. **b,** Example units as in **a** but for sessions with 8 start positions rather than 4. **c,** The spatial sparsity of units in 4-start sessions (0.33, [0.30, 0.37], n=206 units) was not significantly different (p=0.56, Wilcoxon rank-sum test) than the spatial sparsity in 8-start sessions (0.35, [0.32, 0.37], n=178 units). **d,** There was no difference in spatial sparsity between 4-start and 8-start positions when controlling for the total number of spikes (p=0.18, two-way ANOVA, see Methods).

**Extended Data Fig. 4.**
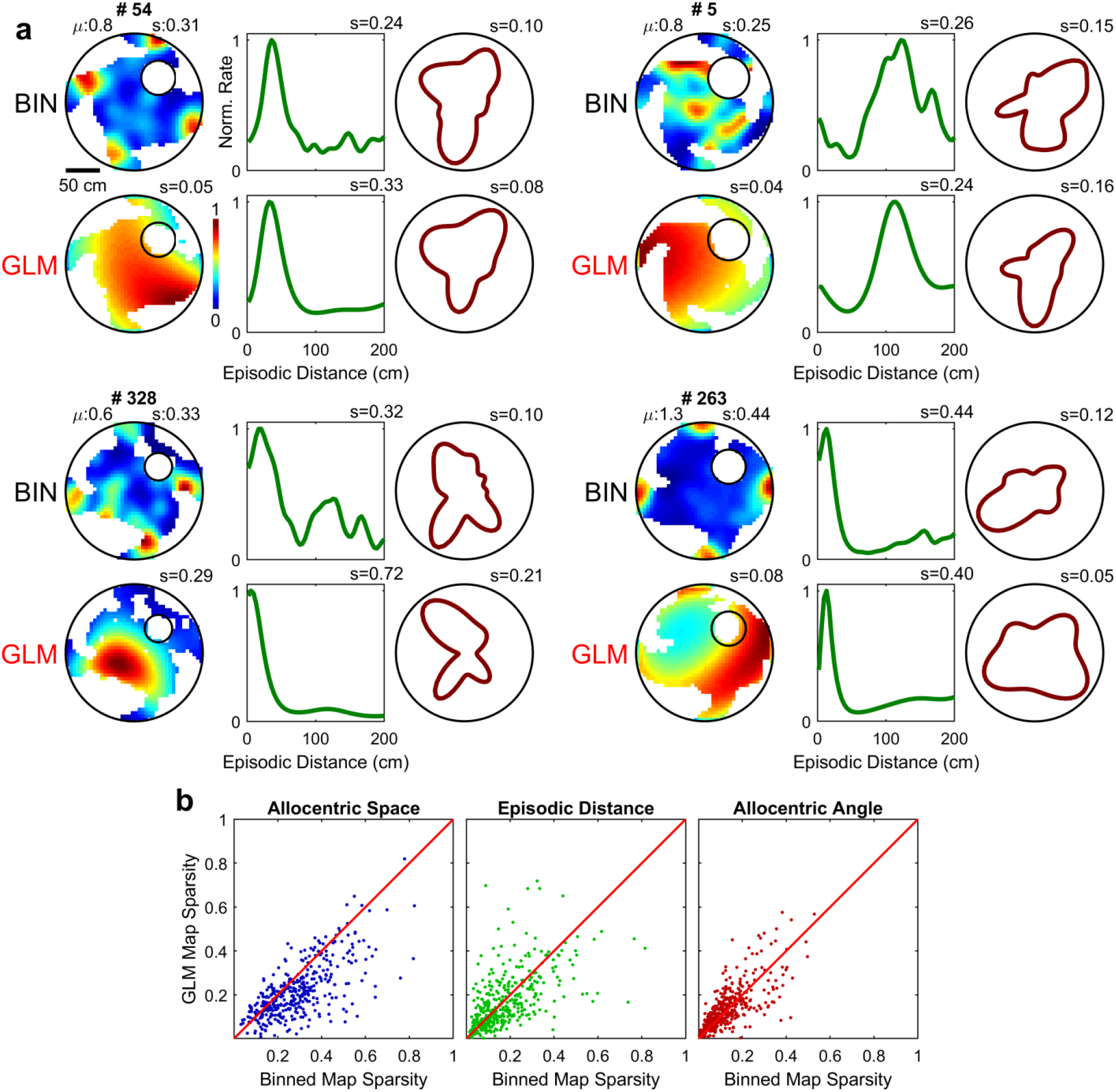
Differences between binning and GLM-derived maps. **a,** 4 example units demonstrating the differences between maps derived from the binning method (top rows) and the GLM (bottom rows). Note that spatial rate maps in particular may seem tuned using the binning method when in fact this can be explained by tuning in either the distance or angular domains. **b,** Sparsity of spatial (left), distance (middle), and angular (right) maps using the binning method versus the sparsity using the GLM. For all variables, the binning method estimated larger sparsity on average than the GLM (Space: p=1.2×10^-59^; Distance: p=1.5×10^-10^; Angle: p=2.7×10^-26^; n=384 units, Wilcoxon sign-rank test for all).

**Extended Data Fig. 5.**
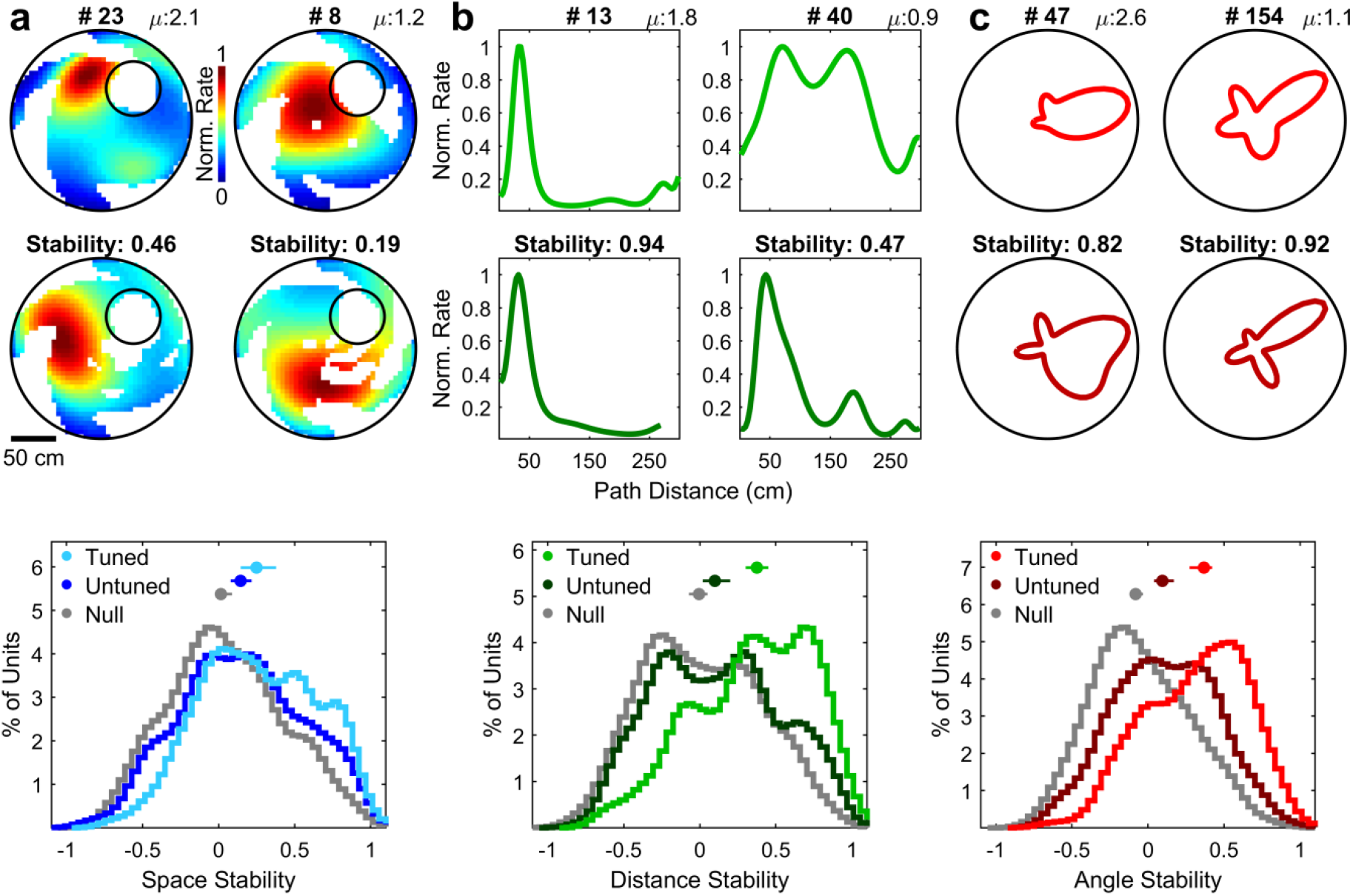
Quantification of stability of GLM results for space, distance, and angle tuning. **a,** Top, example rate maps for two units from the first half (top row) and second half (middle row) of a session demonstrating different levels of stability. Bottom, the stability of tuned spatial rate maps (0.25, [0.14, 0.40], n=111 units) was significantly higher than both the stability of untuned maps (0.14, [0.08, 0.23], n=273 units; p=0.02, Wilcoxon rank-sum test here and throughout the figure) and the stability expected from random shuffles of first and second half maps (−0.00, [-0.06 0.08], n=384 units; p=3.7×10^-7^). Untuned maps were also more stable than chance (p=5.2×10^-4^). **b,** Example path distance rate maps for two units from the first half (top row) and second half (bottom row) of a session demonstrating different levels of stability. Bottom, the stability of tuned path distance maps (0.38, [0.30 0.44], n=181 units) was significantly higher than the stability of untuned maps (0.10, [0.03 0.20], n=203 units; p=5.1×10^-8^) and of shuffled controls (−0.02, [-0.10 0.03], n=384 units; p=1.8×10^-19^). Untuned distance maps were also more stable than chance (p=3.2×10^-3^). **c,** Example angle rate maps for two units from the first half (top row) and second half (middle row) of a session demonstrating different levels of stability. Bottom, the stability of tuned path angle maps (0.37, [0.28 0.43], n=155 units) was significantly higher than the stability of untuned maps (0.09, [0.05 0.17], n=229 units; p=1.9×10^-8^) and of shuffled controls (0.02, [-0.04 0.05], n=384 units; p=3.4×10^-17^). Untuned distance maps were also more stable than chance (p=1.7×10^-3^).

**Extended Data Fig. 6.**
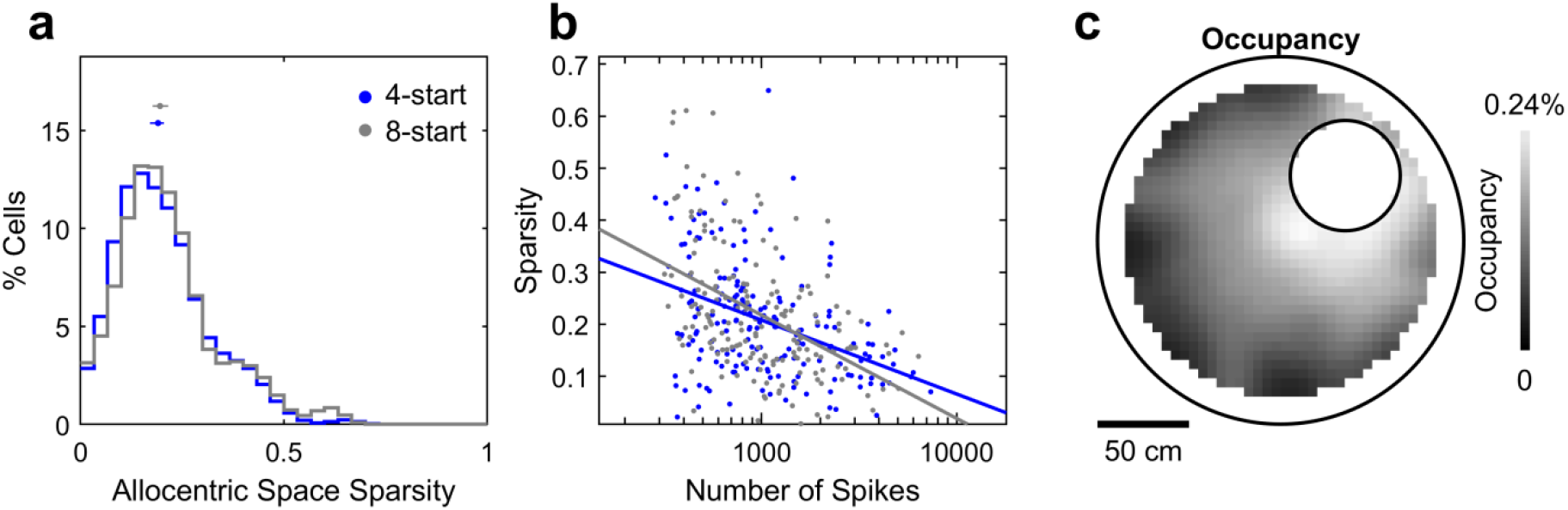
GLM-derived spatial sparsity and spatial occupancy. **a,** The spatial sparsity of units in 4-start sessions (0.33, [0.30, 0.37], n=206 units) was not significantly different (p=0.56, Wilcoxon rank-sum test) than the spatial sparsity in 8-start sessions (0.35, [0.32, 0.37], n=178 units). **b,** There was no difference in spatial sparsity between 4-start and 8-start positions when controlling for the total number of spikes (p=0.18, two-way ANOVA, see Methods). **c,** The distribution of spatial occupancy averaged across all sessions was clustered towards the goal location, mirroring the pattern seen in the clustering of spatial field peaks (Fig. 2e).

**Extended Data Fig. 7.**
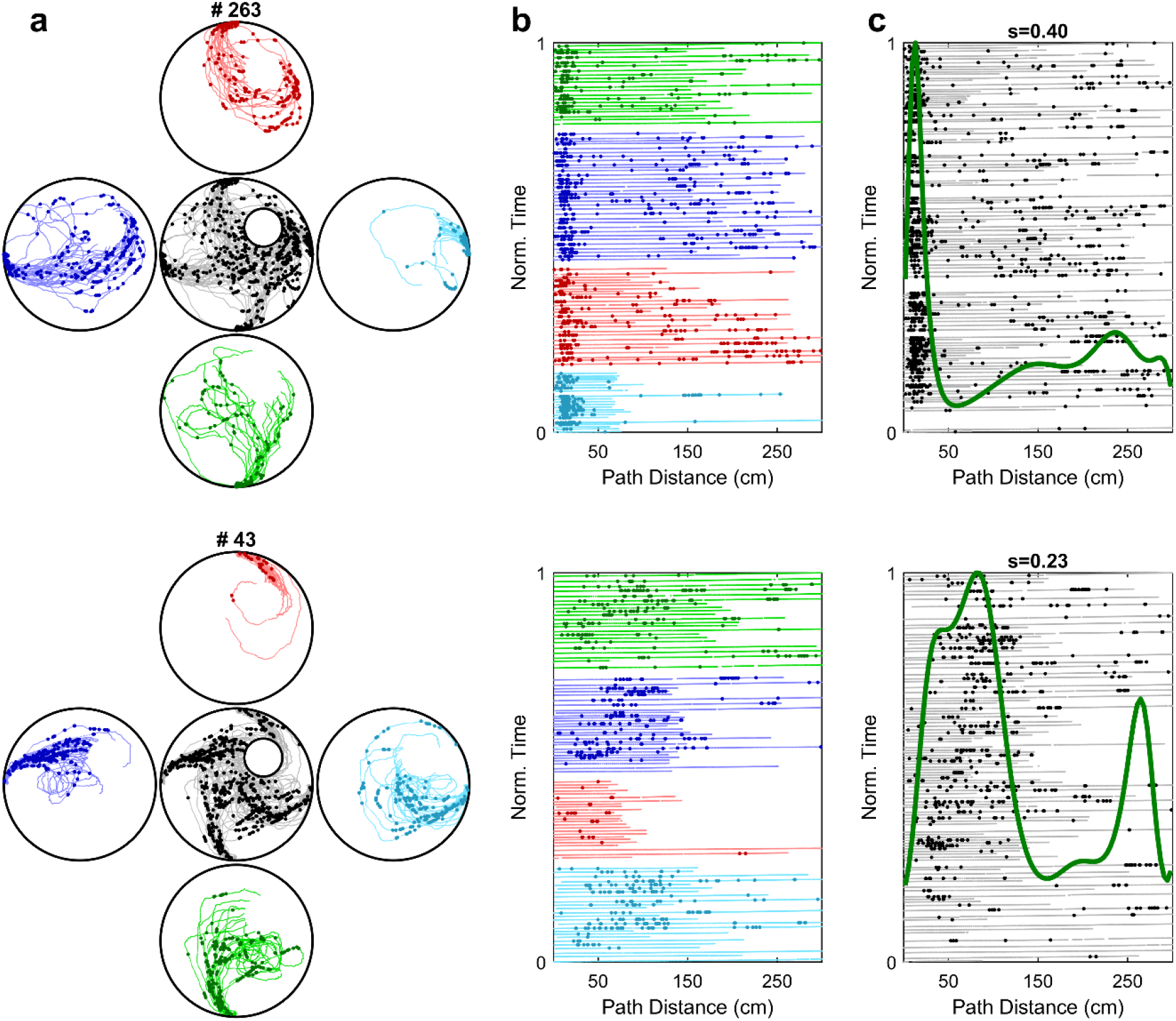
Distance coding cells show similar selectivity across start positions. **a,** Spikes as a function of rat’s position, for two different cells (top and bottom) are color coded based on the start position. **b,** Spikes as a function of the distance traveled, with trials from different start positions grouped together. The maps look qualitatively similar from all four start positions. The variations in firing rates could occur due to other variables, e.g. direction selectivity. **c,** Hence, we used the GLM method (see Methods) using data from all the trials. Spikes are shown as a function of the path distance and time elapsed. The GLM estimate of firing rate as a function of distance alone is shown by thick line.

**Extended Data Fig. 8.**
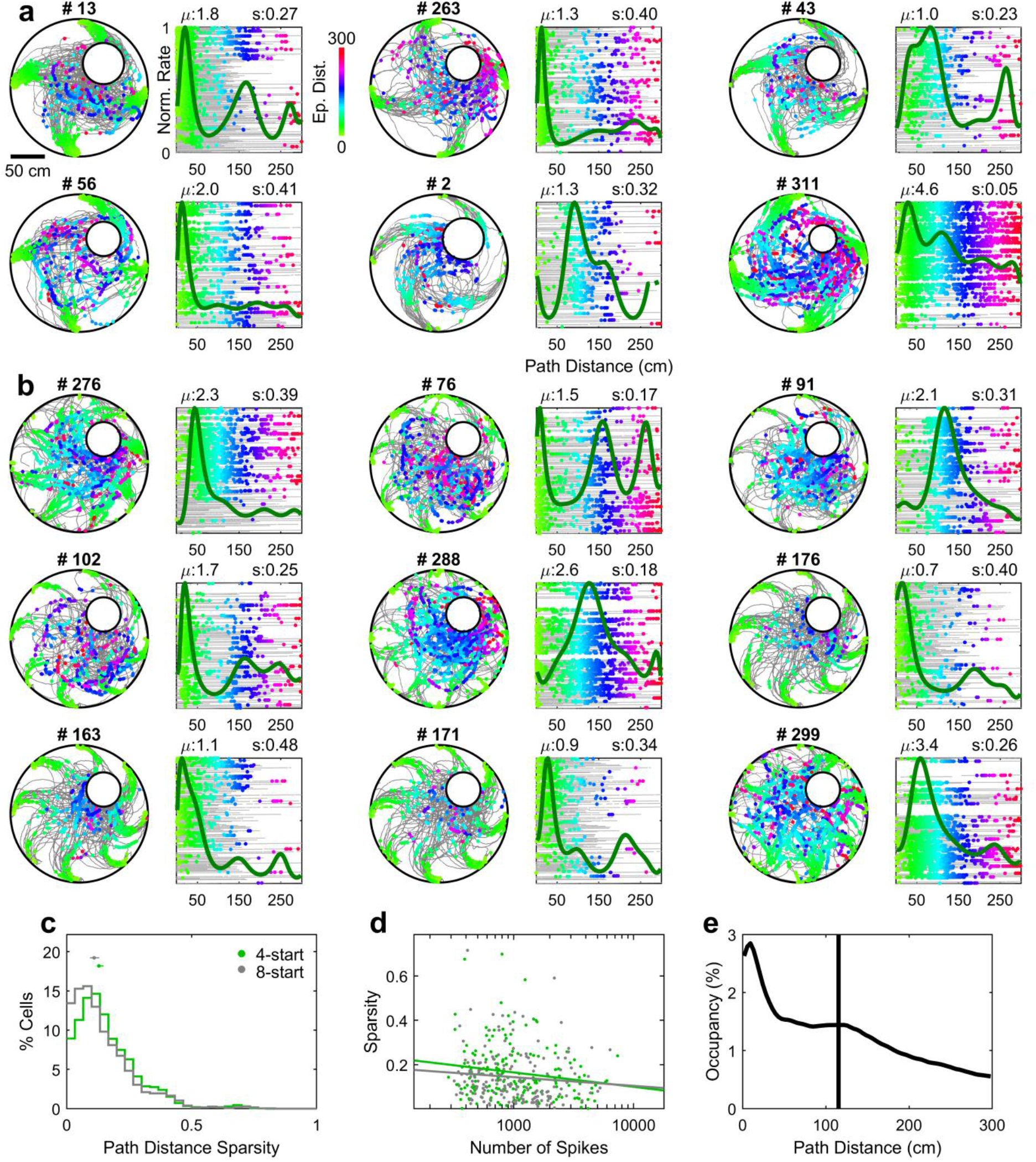
Examples of path distance tuning for longer distances in 4- and 8-start navigation tasks. **a,** Example units as in Fig. 2b. **b,** Example units as in **a** but for sessions with 8 start positions rather than 4. **c,** The distance sparsity of units in 4-start sessions (0.13, [0.12, 0.14], n=183 units) was slightly but significantly greater I (p=0.03, Wilcoxon rank-sum test) than the distance sparsity in 8-start sessions (0.11, [0.09, 0.13], n=181 units). **d,** The effect in **c** was not present when controlling for the total number of spikes (p=0.43, two-way ANOVA, see Methods). **e,** The distribution of occupancy times was skewed toward earlier distances, with a center of mass at 115 cm.

**Extended Data Fig. 9.**
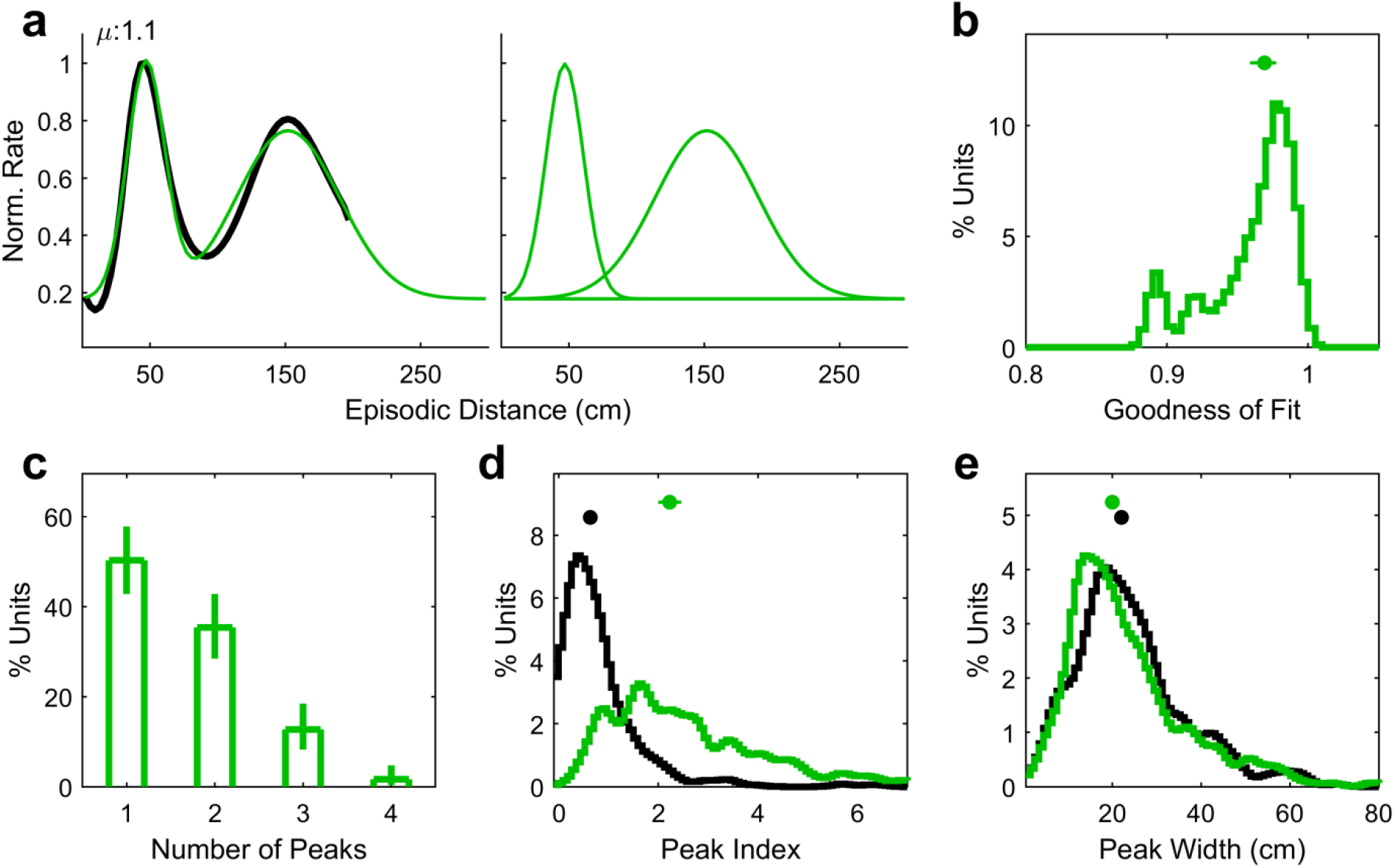
Additional properties of path distance tuning. **a,** Left, sample distance tuning curve (black) overlaid with the sum of two fitted Gaussians (green). Right, the individual Gaussians that were fitted. **b,** The median goodness of fit (correlation coefficient between the original and fitted curve) was quite high (0.97, [0.96, 0.98], n=181 units), with no unit having a fit less than 0.89. **c,** Distribution of the number of significant peaks in distance maps. 50% of units had more than one peaks, with a mean of 1.7, [1.5, 1.8] peaks. Error bars represent the 95% confidence interval of the mean obtained from a binomial distribution using the Matlab function *binofit()*. **d,** The peak index (peak amplitude of a fitted Gaussian divided by constant offset) of distance curves (2.2, [2.0, 2.4], n=300 peaks) was significantly higher (p=2.1×10^-69^, Wilcoxon rank-sum test here and in **e**) than for shuffled data (0.63, [0.57, 0.68], n=463 peaks). **e,** The width of fitted Gaussian components (width at half-max; 20, [18, 21] cm, n=300 peaks) was slightly but significantly smaller (p=0.03) than for shuffled data (22, [21, 23] cm, n=463 peaks).

**Extended Data Fig. 10.**
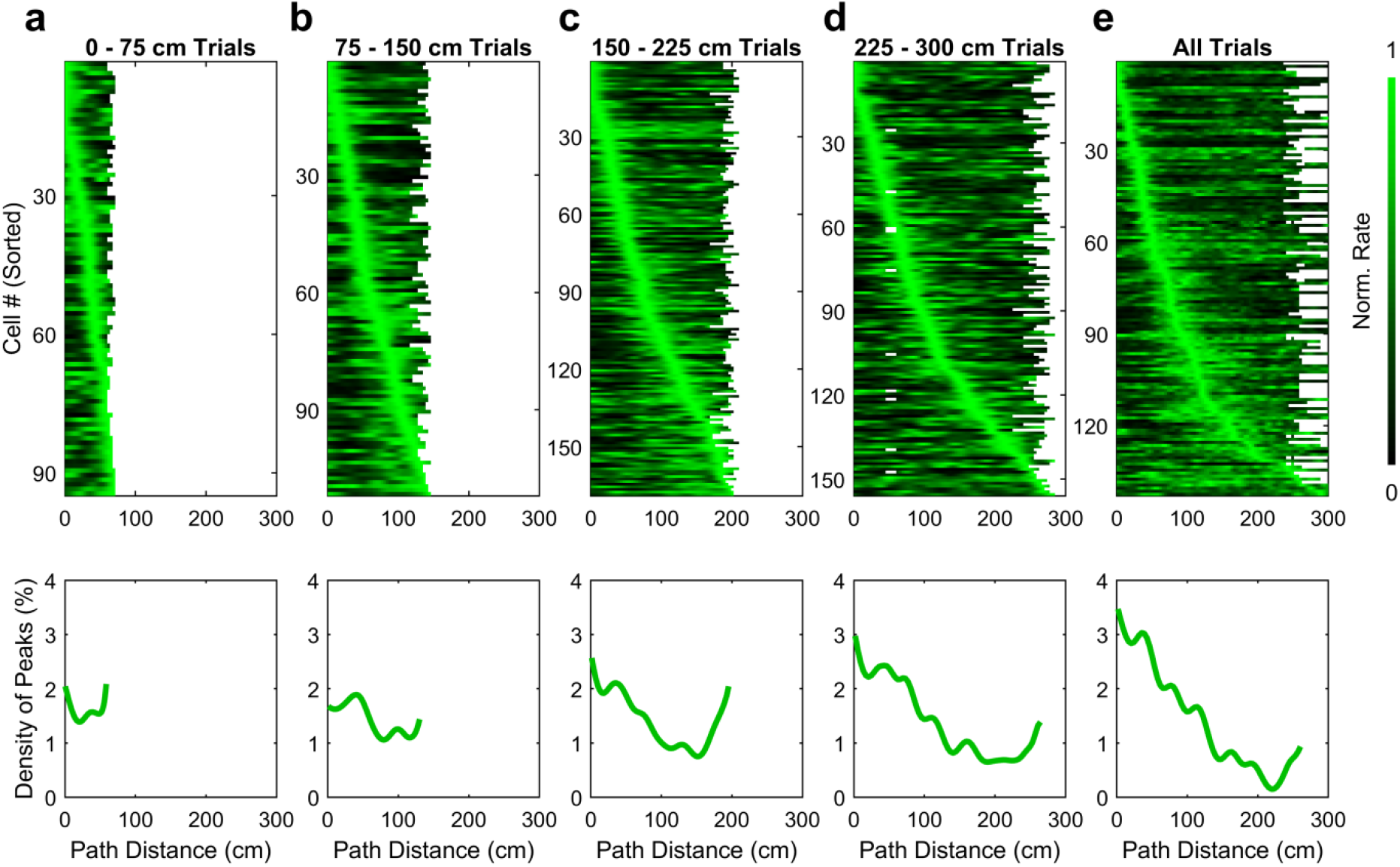
Path distance centers are aggregated towards short distances independent of trial length. **a)** Top, binned distance rate maps for all cells significantly tuned for path distance, sorted by location of peak rate, using only data from trials of length 0 – 75 cm (see Methods for cell inclusion criteria). Bottom, distribution of the peak location of rate maps above (Median of 36, [32, 41] cm, n=94 peaks). **b)** Same as **a** but computed using only data from trials of length 75 – 150 cm (Median of 54, [43, 69] cm, n=111 peaks). **c)** Same as a but computed using only data from trials of length 150 – 225 cm. Note there is still aggregation of field peaks near the beginning of paths (Median of 71, [54, 81] cm, n=168 peaks), even though all paths in this data covered at least 150 cm. **d)** Same as **a**, but for the longest trials, of length 225 – 300 cm. The aggregation towards the beginning of trials (Median of 77, [69, 96] cm, n=155 peaks) is quite pronounced. **e)** Same as panels **a-d** but including data from all trials (Median of 71, [51, 84] cm, n=142 peaks). Note that these maps are computed using the binning method, and thus differ slightly from those in Fig. 2f which are computed using the GLM.

**Extended Data Fig. 11.**
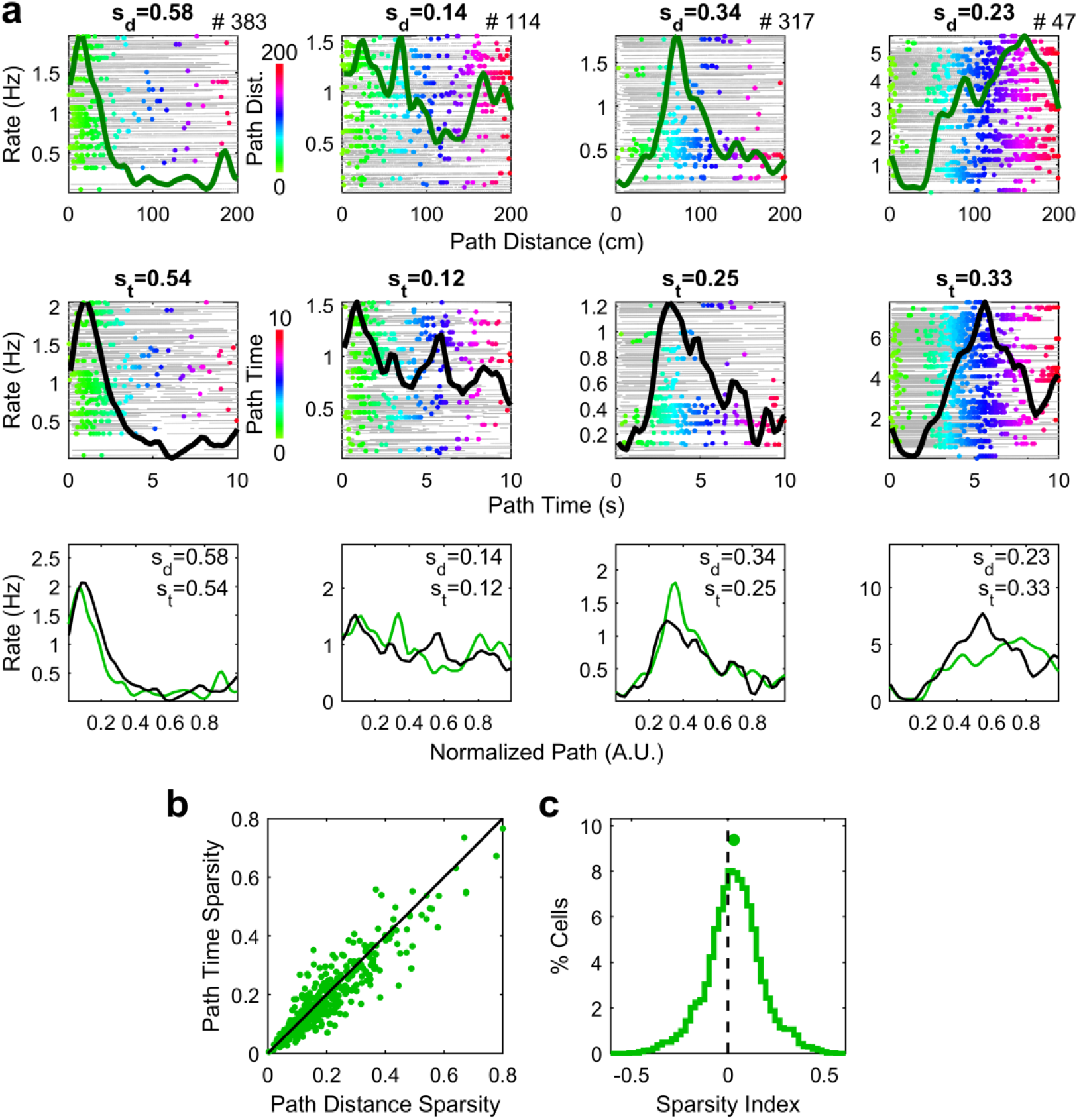
Comparison of selectivity for distance versus time. **a,** Path distance (top row) and path time (middle row) rate maps for four sample cells. s_d_ and s_t_ represent the sparsity of rate maps for distance and time, respectively. Rate maps are overlaid in the bottom row for ease of comparison. Distance between 0 and 200 cm and time between 0 and 10 seconds are normalized from 0 to 1 for visualization. Column 1 depicts a cell that is well-tuned in both the distance and time domains. Column 2 depicts a cell that is poorly-tuned in both domains. Column 3 shows a cell that is better tuned in the distance domain. Column 4 shows a cell that is better tuned in the time domain. **b,** Sparsity of Path Time maps versus sparsity of Path Distance maps. **c**, Sparsity index (defined as (s_d_ – s_t_)/(s_d_ + s_t_)) was slightly but significantly greater than 0 (0.03, [0.02 0.05], n=384 cells; p=1.5×10^-6^, Wilcoxon sign-rank test).

**Extended Data Fig. 12.**
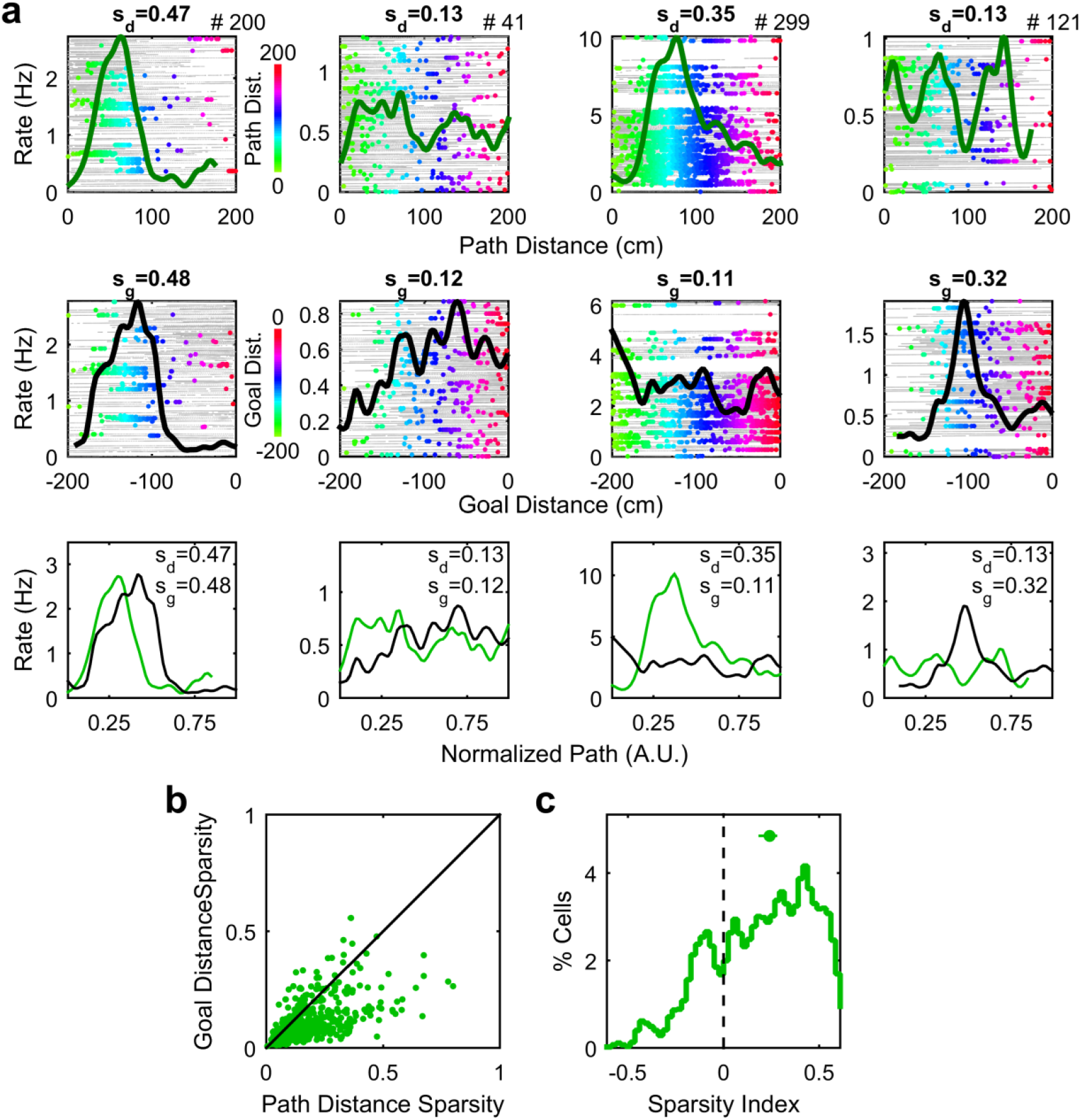
Neurons encode distance significantly better with respect to the start position than the goal position. **a,** Path distance (top row) and goal distance (middle row) rate maps for four sample cells. s_d_ and s_g_ represent the sparsity of rate maps for path distance and goal distance, respectively. Rate maps are overlaid in the bottom row for ease of comparison. Distance between 0 and 200 cm and time between −200 and 0 cm are normalized from 0 to 1 for visualization. Column 1 depicts a cell that is well-tuned in both the frames of reference. Column 2 depicts a cell that is poorly-tuned in both domains. Column 3 shows a cell that is better tuned in the path distance frame. Column 4 shows a cell that is better tuned in the goal distance domain. **b,** Sparsity of Goal Distance maps versus sparsity of Path Distance maps. **c**, Sparsity index (defined as (s_d_ – s_g_)/(s_d_ + s_g_)) was significantly greater than 0 (0.27, [0.23 0.31], n=384 cells; p=7.8×10^-37^, Wilcoxon sign-rank test).

**Extended Data Fig. 13.**
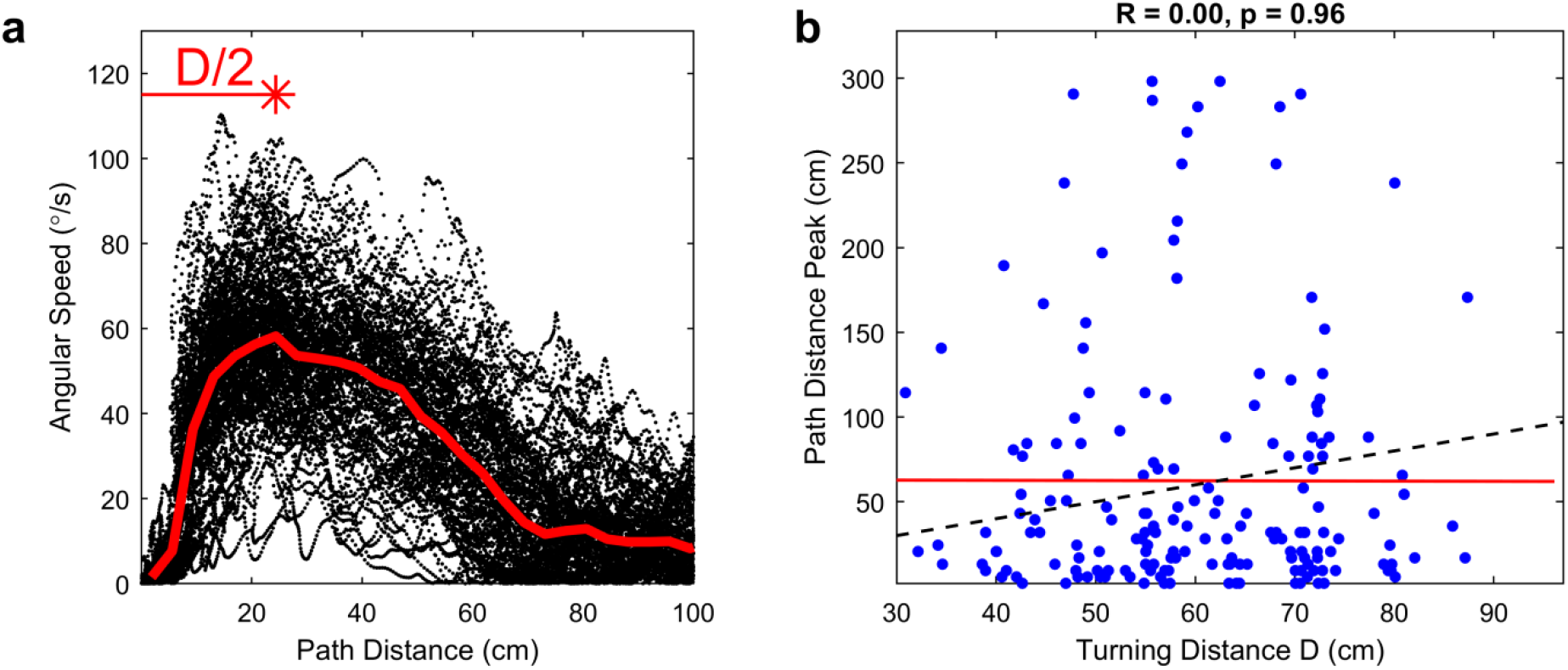
Path distance coding does not depend on turning distance. **a,** Example calculation of turning distance D. Path Distance versus angular speed shows the repeated movement trajectory across trials (black dots). The thick red line is the median angular speed as a function of path distance (computed in bins of width 3.75 cm, smoothed with a Gaussian kernel with a sigma of 3.75 cm). The distance corresponding to the peak angular speed (red star) indicates the halfway point of the turning distance, or D/2, for that session. **b)** For each cell significantly tuned for path distance, the location of the peak is plotted against the turning distance D for the corresponding session. There is no correlation between the two (R = 0.00, p = 0.96, two-sided *t* test, n=176 cells), indicating that path distance peaks are not solely defined by turning. The dotted black line indicates the unity line. For many cells, the path distance peak was substantially larger than the turning distance, additionally indicating that distance selectivity was not entirely determined by the act of turning. For ease of viewing, random Gaussian jitter (sigma of 2 cm) is added to the turning distance for each cell. Statistics and the red best fit line are computed on the original data with no jitter. 5 cells were excluded from the original 181 distance-tuned cells for belonging to sessions with a turning radius > 150 cm.

**Extended Data Fig. 14.**
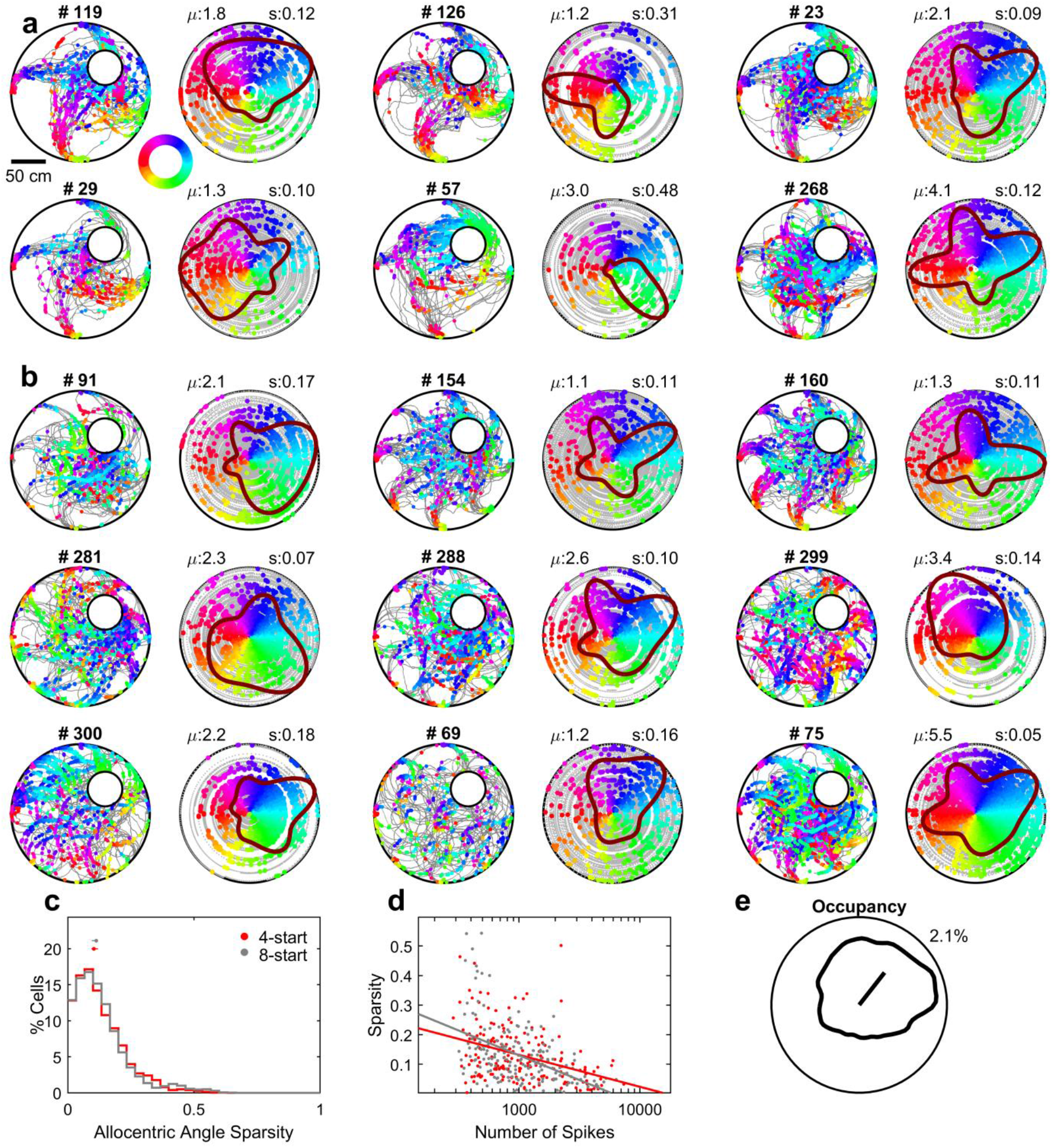
Additional examples of angular tuning in 4- and 8-start navigation tasks. **a,** Example units as in Fig. 2c. **b,** Example units as in a but for sessions with 8 start positions rather than 4. **c,** The angular sparsity of units in 4-start sessions (0.10, [0.09, 0.12], n=155 units) was not significantly different (p=0.77, Wilcoxon rank-sum test) than the angular sparsity in 8-start sessions (0.11, [0.09, 0.12], n=155 units). **d,** There was no significant difference when controlling for the total number of spikes (p=0.06, two-way ANOVA, see Methods). **e,** The distribution of occupancy times was skewed toward the north-east direction, with a mean vector pointing towards 56°.

**Extended Data Fig. 15.**
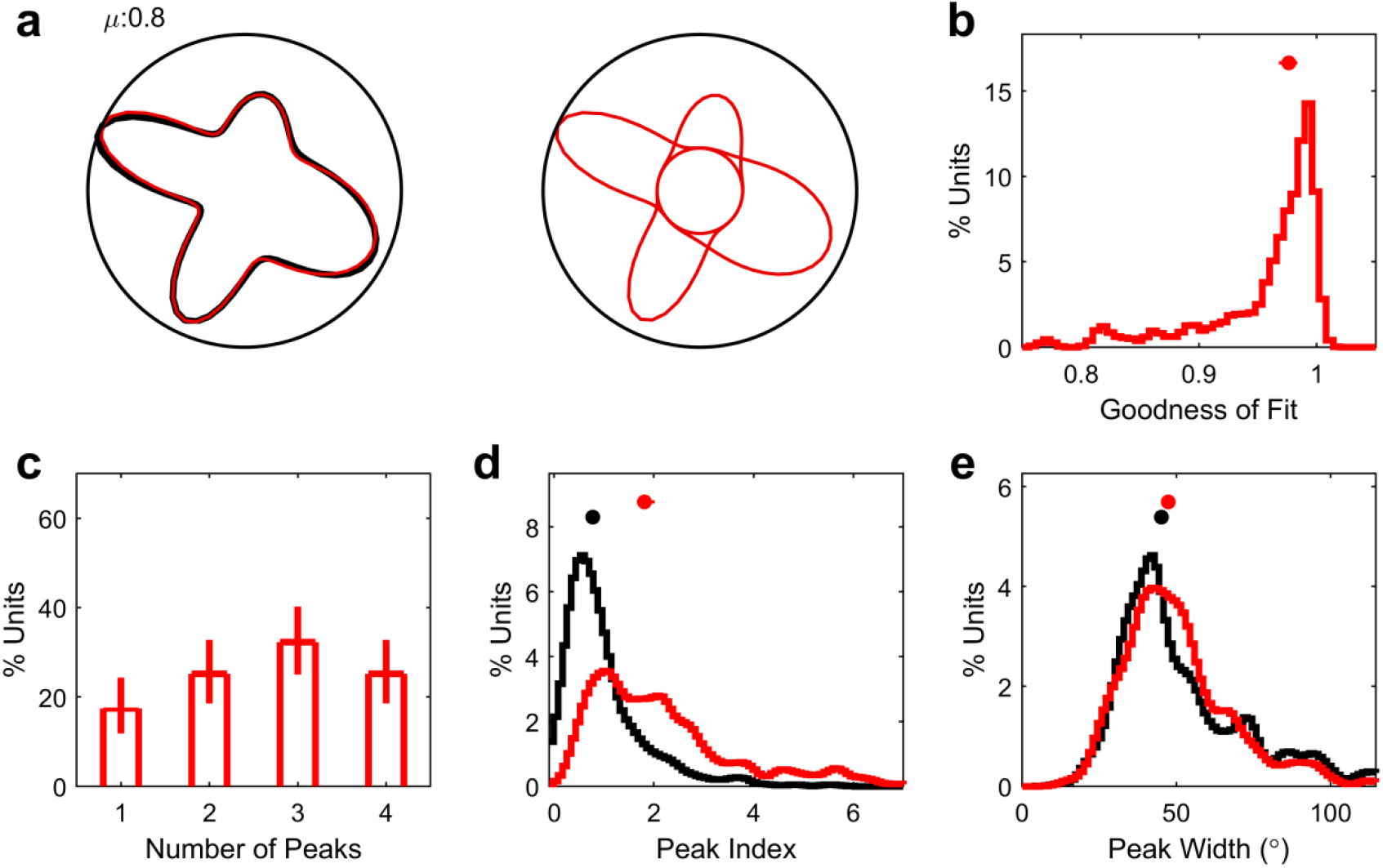
Additional properties of angular tuning. **a,** Left, sample angle tuning curve (black) overlaid with the sum of four fitted Von Mises curves (red). Right, the individual Von Mises curves that were fitted. **b,** The median goodness of fit (correlation coefficient between the original and fitted curve) was quite high (0.98, [0.97, 0.98], n=155 units). **c,** Distribution of the number of significant peaks in angle maps. 83% of units had more than one peak, with a mean of 2.7, [2.5, 2.8] peaks. Error bars represent the 95% confidence interval of the mean obtained from the Matlab function *binofit()*. **d,** The peak index (peak amplitude of a fitted Von Mises curve divided by constant offset; 1.8, [1.7, 2.0], n=411 peaks) was significantly higher (p=2.1×10^-35^) than for shuffled data (0.77 [0.72, 0.84], n=476 peaks). **e,** The width of fitted Von Mises curves (width at half-max; 47, [46, 49]°, n=411 peaks) was not significantly different (p=0.70) than for shuffled data (45, [44, 48]°, n=476 peaks).

**Extended Data Fig. 16.**
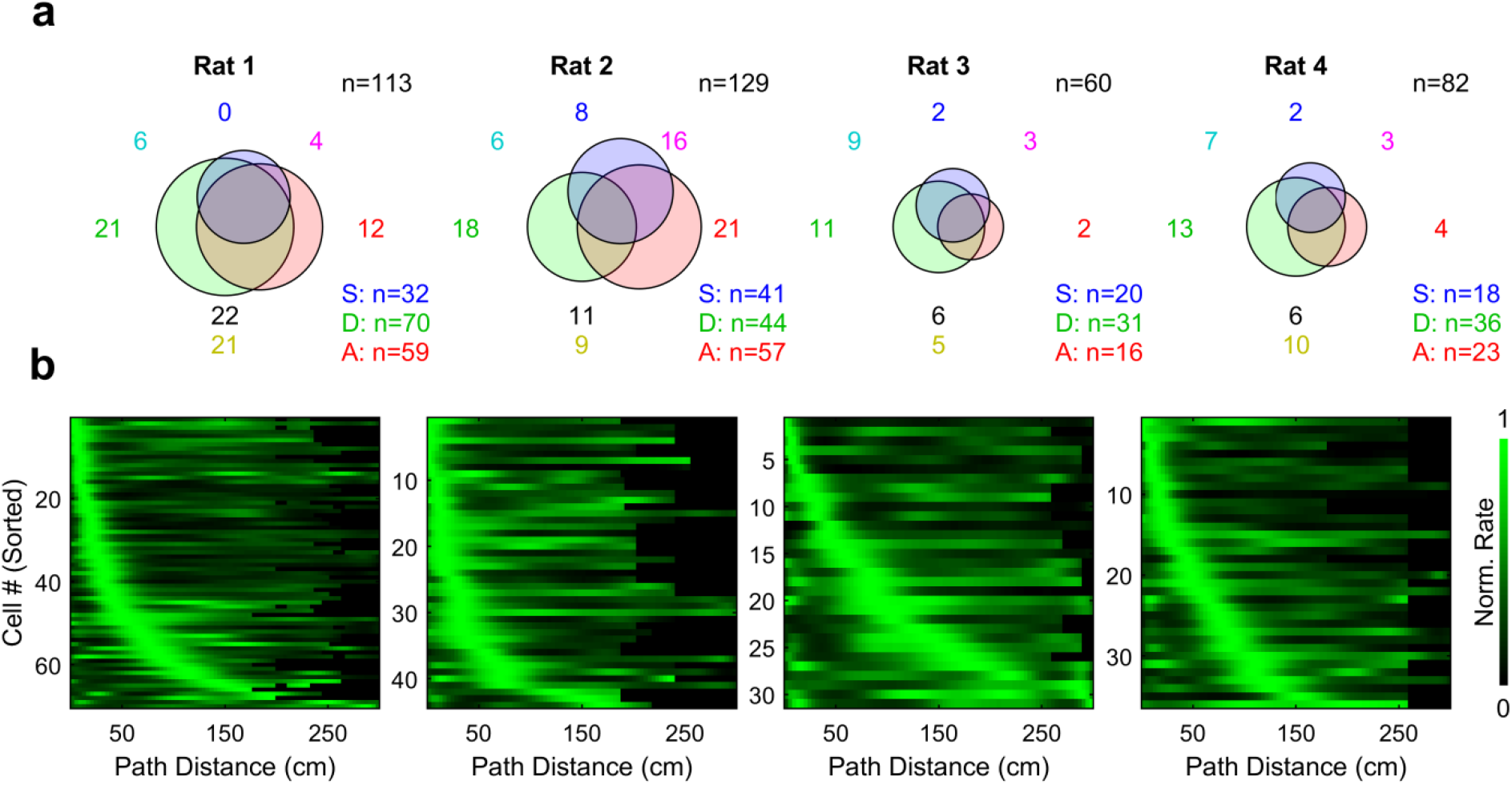
Population tuning measures and distribution of distance fields for individual rats. **a,** As in Fig. 2b for individual rats. The venn diagrams represent the number of cells significantly tuned for Allocentric Space (S, blue), Path Distance (D, green), or Allocentric Angle (A, red). The colored numbers represent the number of cells falling into each region of the venn diagram (Blue, Space only; Green, Distance only; Red, Angle only; Cyan, Space and Distance; Magenta, Space and Angle; Yellow, Distance and Angle; Black, Space, Distance, and Angle). **b,** As in Fig. 2f for individual rats, showing qualitatively similar distributions of distance fields.

**Extended Data Fig. 17.**
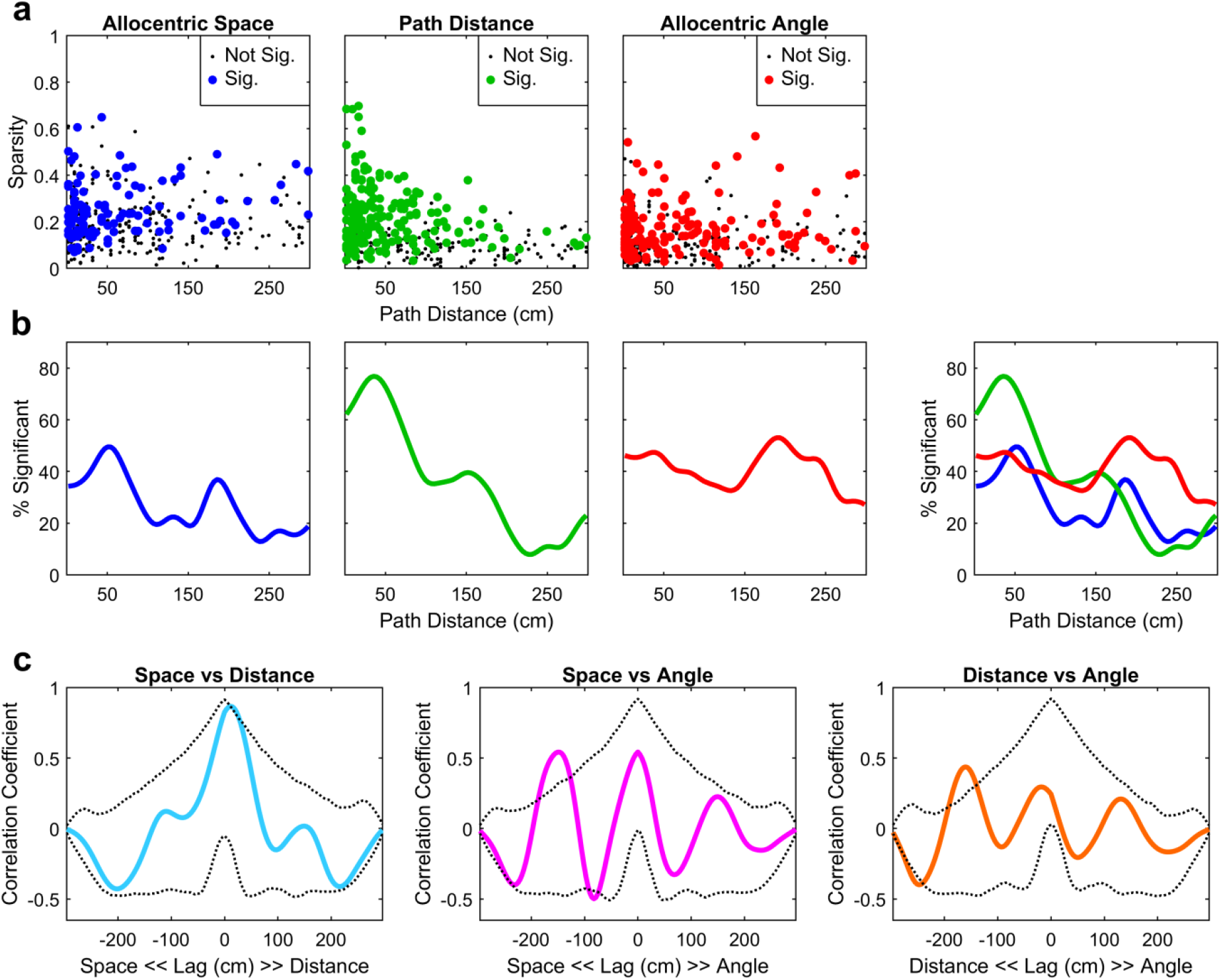
Episodic distribution of space, distance, and angle selectivity. **a,** Sparsity for rate maps in allocentric space (left), path distance (middle), and allocentric angle (right) versus the center distance coordinate (see Methods) for each cell. Significantly tuned cells are marked with large, colored dots and cells that are not significantly tuned are marked with small, black dots. **b,** The percentage of cells significantly tuned as a function of their center distance coordinate for space (blue), distance (green), and angle (red). The combined plot at the far right is the same as Fig. 2h. **c,** Cross-correlations between the curves in **b,** overlaid with shuffled control cross-correlations, demonstrate that the relative ordering of parameter tuning – Distance, then Space, then Angle – is greater than expected by chance. Dotted black lines indicate the median and 95% range of the cross-correlation of the curves in **b** constructed from shuffled data (see Methods). Cross-correlation peaks above this range (Left, cyan, 11.25 cm indicating Distance leads Space; Middle, magenta, −150 cm indicating Space leads Angle; Right, orange, −161.3 cm indicating Distance leads Angle) indicate statistical significance at the p<0.05 level.

**Extended Data Fig. 18.**
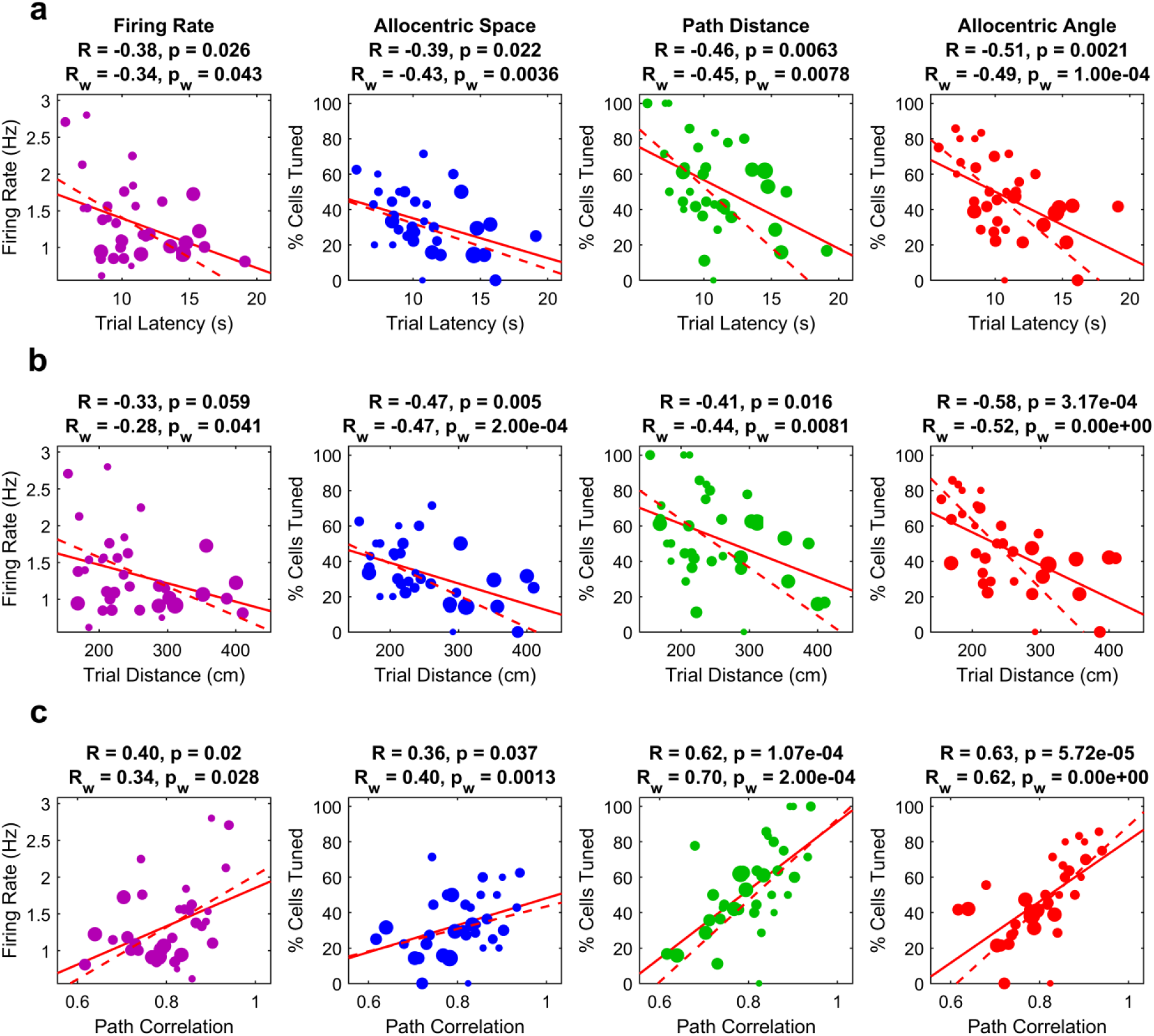
Additional measures of performance correlate with neural tuning. **a,** Same values as in Fig. 4c plotted as a function of Trial Latency (see Methods). Correlation values and p-values are shown above each figure. R_w_ and p_w_ represent the correlation value and p-value for the weighted best fit line. For unweighted fits, p-values are from a two-sided *t* test for each panel, with n=34 sessions. For weighted fits, p-values are calculated through a resampling procedure (see Methods). **b,** Same as **a**, but plotted as a function of Trial Distance (see Methods). **c,** Same as **a**, but plotted as a function of within-start path correlation (Extended Data Fig. 1, see Methods).

**Extended Data Fig. 19.**
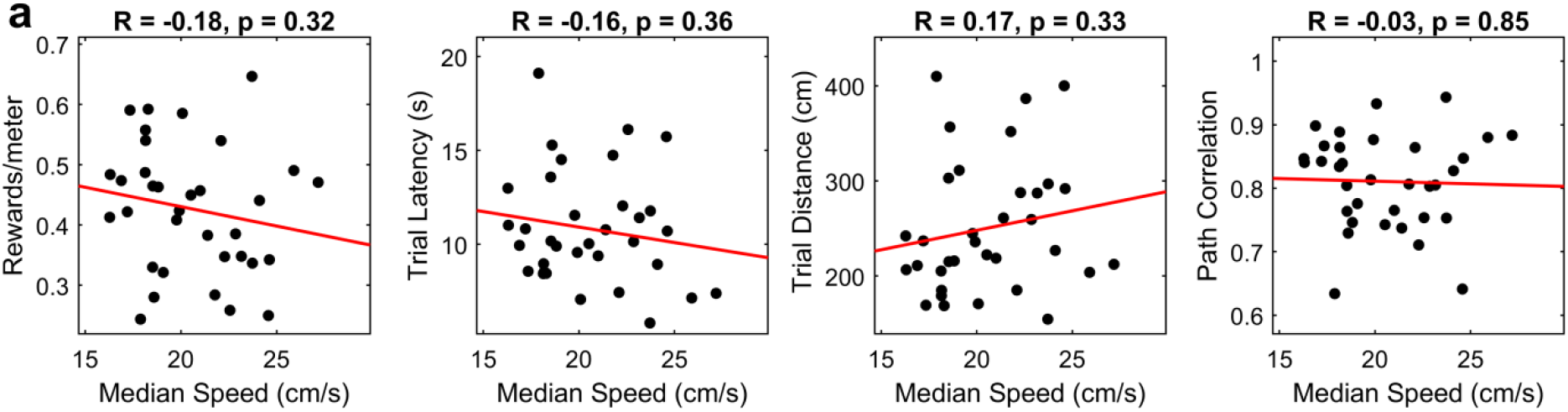
Speed does not correlate with performance. **a,** Including sessions from all rats, the median speed in a session was not significantly correlated with behavioral performance as measured by rewards/meter, Trial Latency, Trial Distance, or Path Correlation. Correlation values and p-values are shown above each figure. p-values are from a two-sided *t* test for each panel, with n=34 sessions.

**Extended Data Fig. 20.**
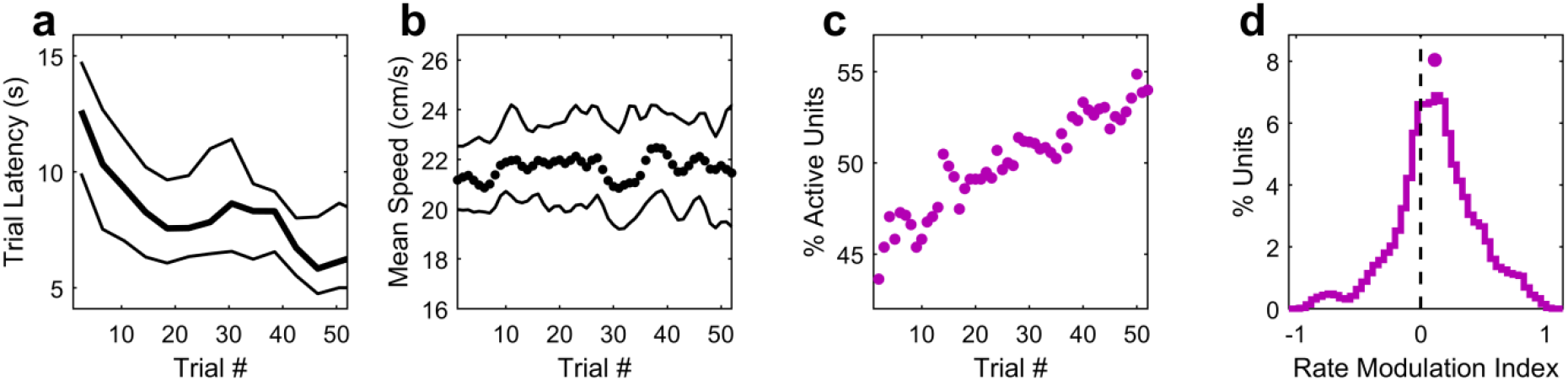
Additional quantification of experience-dependent changes in behavior and neural activation. **a,** Trial latency (time spent running) decreased as a function of trial number (Effect of trial number on Trial Latency: p=5.0×10^-5^, 34 sessions; one-way repeated-measures ANOVA). The thick line is the median, and the thin lines are the 95% confidence interval of the median. **b,** Mean speed did not significantly change as a function of trial number (Effect of trial number on Mean Speed: p=0.44, 34 sessions; one-way repeated-measures ANOVA). The thick line is the median, and the thin lines are the 95% confidence interval of the median. **c,** The fraction of total cells active (rate in a 3-trial boxcar average > 0.2 Hz) increased with experience (correlation coefficient R=0.94, p=2.7×10^-25^, n=52 trials, two-sided *t* test). **d,** Averaged across the ensemble, the firing rates were higher in later trials than early trials (Figure 4b). This was quantified on a cell-by-cell basis by computing the rate modulation index for each cell (see Methods), which was centered at 0.11, [0.08, 0.15], and significantly greater than 0 (p=1.8×10^-13^, Wilcoxon sign-rank test, n=384 units).

**Extended Data Fig. 21.**
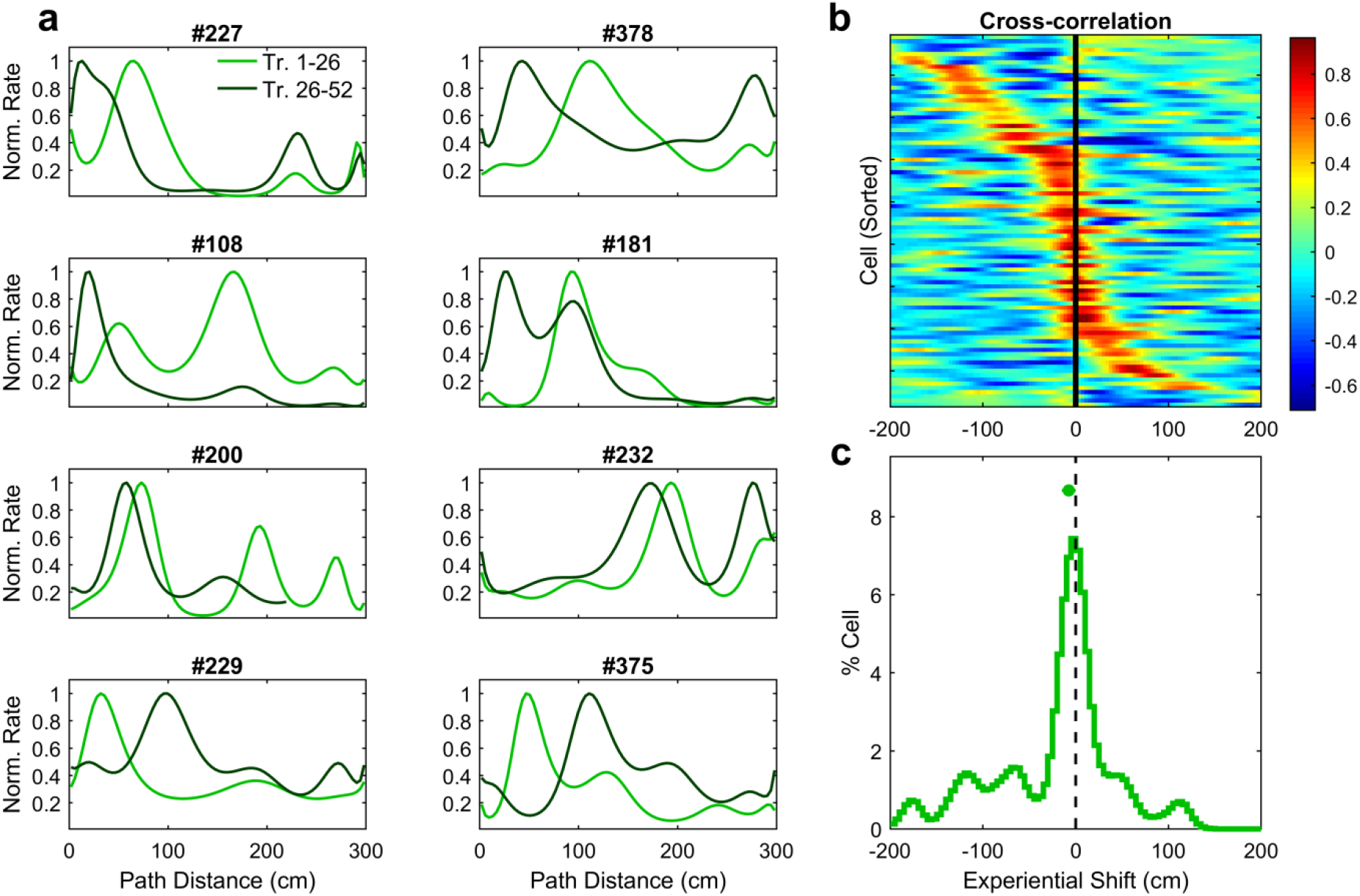
Experience-dependent shifts of path distance tuning in single units. **a,** Eight example cells with distance tuning curves exhibiting shifting with experience. Curves are estimated from the GLM using data only from trials 1-26 (light green) or 27-52 (dark green). The top three rows demonstrate backwards, or anticipatory, shifting. The bottom row demonstrates forwards shifting. **b,** Cross-correlation plots, sorted by the experiential shift (distance lag of the peak in the cross-correlation) of peak correlation, for all cells significantly tuned for distance but not angle, (n=88; 91 cells met this criteria, but 3 were excluded for having insufficient spiking in one of the two halves). c, The median experiential shift (−7.5, [-15, 0] cm) was significantly different from 0 (p=0.014, Wilcoxon sign-rank test).

**Extended Data Fig. 22.**
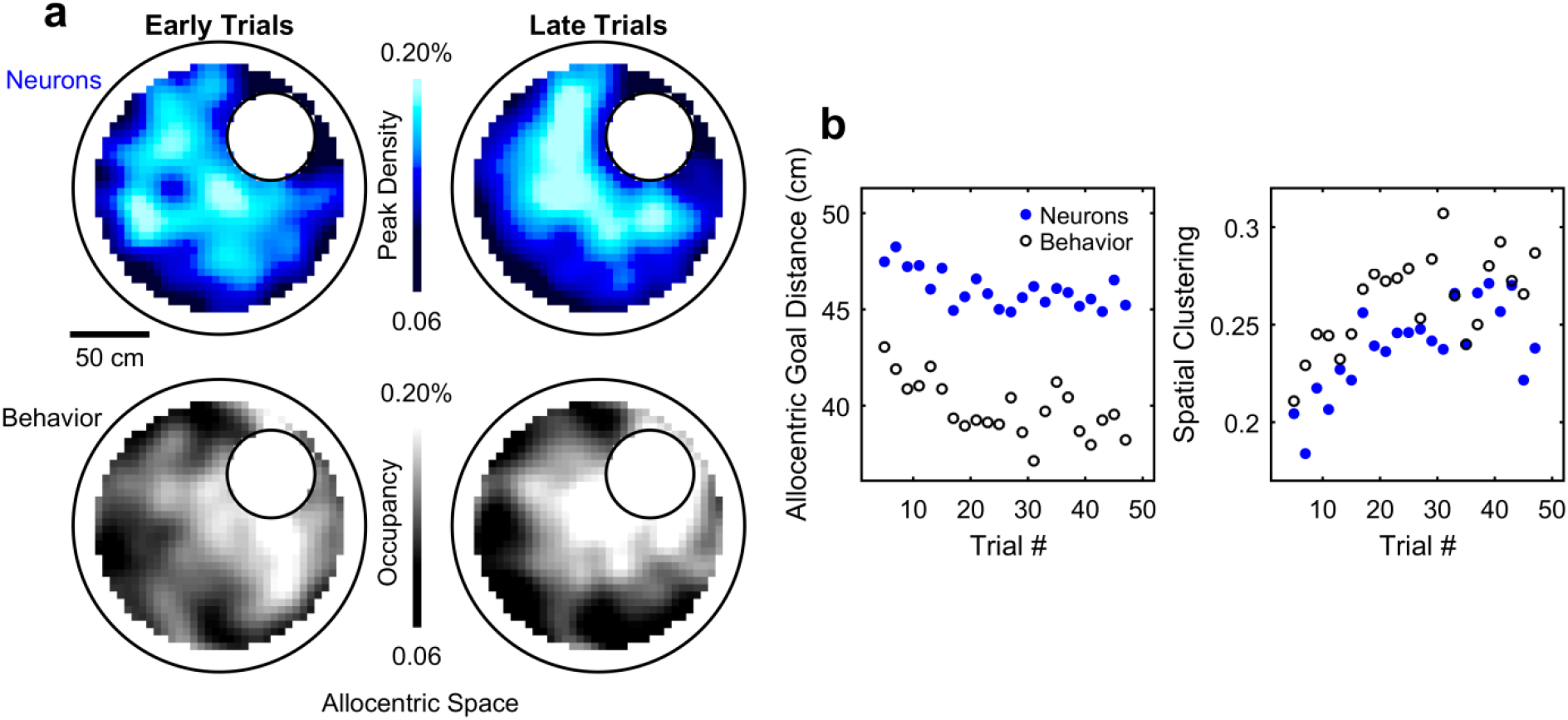
Within-session clustering and forward movement of allocentric spatial maps and their relationship with psychometric curves. **a,** Left, distributions of peaks of allocentric spatial rate maps (top, blue) and spatial occupancy (bottom, gray) in early trials, showing dispersed, fairly uniform distributions. Right, distributions of spatial peaks (top) and spatial occupancy (bottom) in later trials, showing clear clustering near the goal location. **b,** The allocentric goal distance (see Methods) significantly decreased with increasing trial number (Neurons: R=-0.63, p=1.5×10^-3^, two-sided *t* test, n=22 trial blocks, defined as in Fig. 4; Behavior: R=-0.64, p=1.2×10^-3^, two-sided *t* test, n=22 trial blocks), and the sparsity of these distributions increased with trial number (Neurons: R=0.67, p=5.7×10^-4^, two-sided *t* test, n=22 trial blocks; Behavior: R=0.63, p=1.8×10^-3^, two-sided *t* test, n=22 trial blocks).

**Extended Data Fig. 23.**
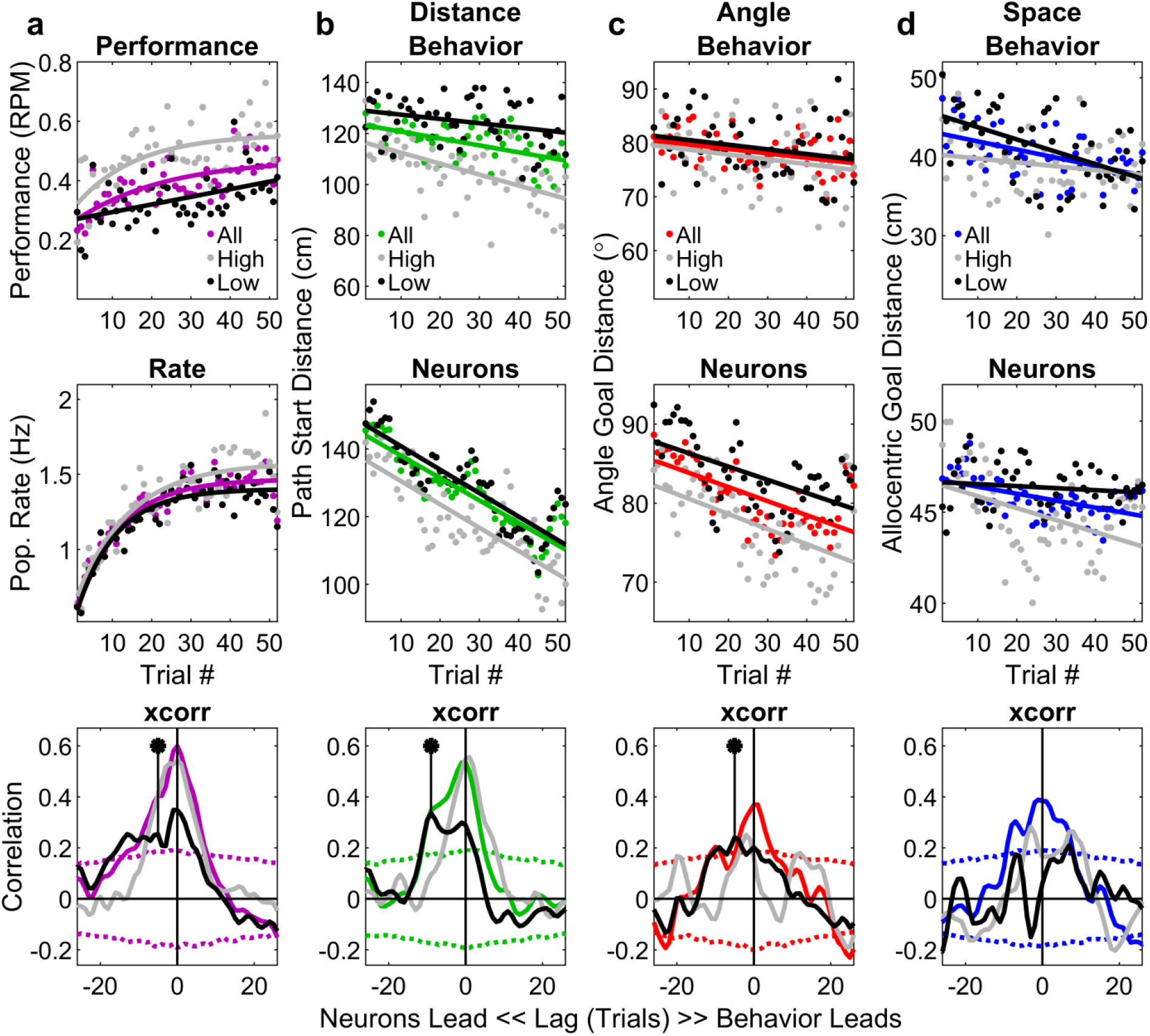
Temporal relationship between neural firing properties and behavior, split into high- and low-performing sessions. **a,** Top, performance increased with trial number. This was true when including all cells (colored dots, same as Fig. 4, right), cells from sessions with high performance (top 50% of sessions, gray dots, “High”), or cells from sessions with low performance (bottom 50% of sessions, black dots, “Low’). Solid lines are exponential fits to the data. Middle, the firing rate of active cells increased with trial number. Bottom, cross-correlation of the population firing rate with performance, for all sessions, high-performance sessions, and low-performance sessions. For all data and high-performance sessions, the lag of the peak correlation is near 0, indicating a co-evolution of performance with firing rate. In low-performance sessions, there is a distinct asymmetry, indicating that neural changes precede behavioral changes. Dotted lines indicate the 99% range of shuffled cross-correlations. The marked point is the approximate center of this asymmetry, at −5 trials, and is above the chance line, indicating statistical significance at the p<0.01 level. **b**, Top, as in Fig. 4d, the center of the distribution of tuning curve peaks shifted towards the trial beginning with experience within a session.

**Extended Data Fig. 24.**
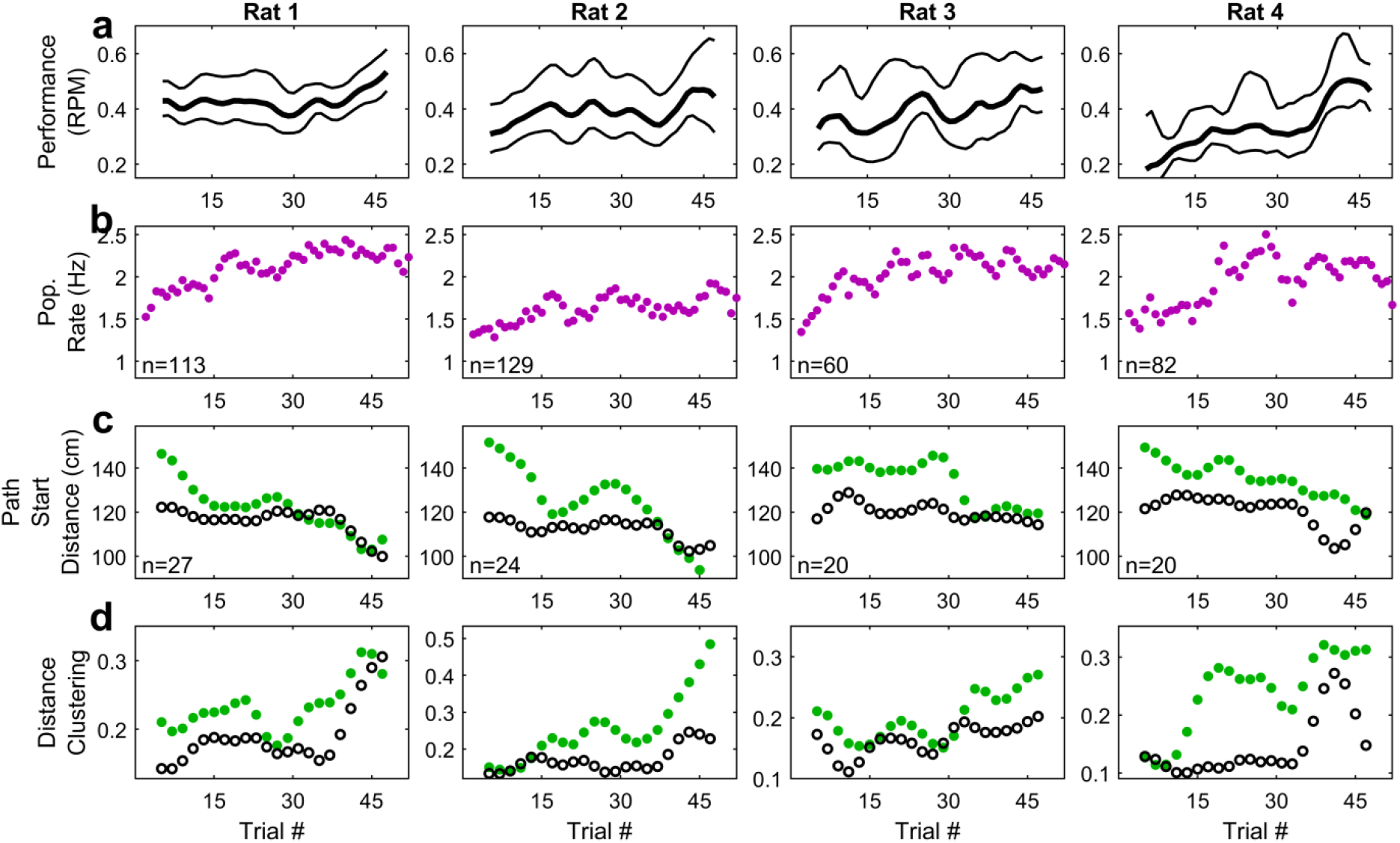
Experience-dependent changes in performance, firing rate, and distance clustering for individual rats. **a,** As in Fig. 4a, but for individual rats, performance increases as a function of trial number (Effect of trial number on performance – Rat 1: p=0.03, 12 sessions; Rat 2: p=0.09, 12 sessions; Rat 3: p=0.11, 6 sessions; Rat 4: p=0.03, 4 sessions; one-way repeated measures ANOVA). **b,** As in Fig. 4b, for individual rats, showing qualitatively similar patterns (Effect of trial number on Population Rate – Rat 1: p=4.1×10^-3^, 113 units; Rat 2: p=0.28, 129 units; Rat 3: p=0.03, 60 units; Rat 4: p=0.04, 82 units; one-way repeated measures ANOVA). **c-d,** As in Fig. 4d, for individual rats, showing qualitatively similar patterns. Green dots represent neural measures, and black circles represent behavioral measures. Correlation coefficients for **c** – Rat 1: Neurons: R=-0.92, p=1.8×10^-9^, two-sided *t* test, n=22 trial blocks (every other trial from trial 5 to 47) here and for all other tests in **c** and **d**; Behavior: R=-0.64, p=1.2×10^-3^; Rat 2: Neurons: R=-0.88, p=9.4×10^-8^; Behavior: R=-0.68, p=4.9×10^-4^; Rat 3: Neurons: R=-0.80, p=8.6×10^-6^; Behavior: R=-0.67, p=7.1×10^-5^; Rat 4: Neurons: R=-0.93, p=3.5×10^-10^; Behavior: R=-0.72, p=1.5×10^-4^. Correlation coefficients for **d** – Rat 1: Neurons: R=-0.68, p=5.7×10^-4^; Behavior: R=0.68, p=4.5×10^-4^; Rat 2: Neurons: R=0.87, p=1.1×10^-7^; Behavior: R=0.67, p=7.0×10^-4^; Rat 3: Neurons: R=0.69, p=3.5×10^-4^; Behavior: R=0.74, p=8.2×10^-5^; Rat 4: Neurons: R=0.82, p=2.5×10^-6^; Behavior: R=0.70, p=3.1×10^-4^.

